# Septin organization is regulated by the Gpa1 Ubiquitination Domain and Endocytic Machinery during the yeast pheromone response

**DOI:** 10.1101/2023.06.16.545321

**Authors:** Cory P. Johnson, Sudati Shrestha, Andrew Hart, Katherine F. Jarvis, Loren E. Genrich, Sarah G. Latario, Nicholas Leclerc, Tetiana Systuk, Matthew Scandura, Remi P. Geohegan, André Khalil, Joshua B. Kelley

## Abstract

The septin cytoskeleton plays a key role in the morphogenesis of the yeast mating projection, forming structures at the base of the projection. The yeast mating response uses the G-protein coupled receptor (GPCR), Ste2, to detect mating pheromone and initiate mating projection morphogenesis. Desensitization of the Gα, Gpa1, by the Regulator of G-protein Signaling (RGS), Sst2, is required for proper septin organization and morphogenesis. We hypothesized that Gpa1 would utilize known septin regulators to control septin organization. We found that single deletions of the septin chaperone Gic1, the Cdc42 GAP Bem3, and the endocytic adaptor proteins Ent1 and Ent2 rescued the polar cap accumulation of septins in the hyperactive Gα. We hypothesized that hyperactive Gα might increase the rate of endocytosis of a pheromone-responsive cargo, thereby altering where septins are localized. Mathematical modeling predicted that changes in endocytosis could explain the septin organizations we find in WT and mutant cells. Our results show that Gpa1-induced disorganization of septins requires clathrin-mediated endocytosis. Both the GPCR and the Gα are known to be internalized by clathrin-mediated endocytosis during the pheromone response. Deletion of the GPCR C-terminus to block internalization partially rescued septin organization. However, deleting the Gpa1 ubiquitination domain required for its endocytosis completely abrogated septin accumulation at the polarity site. Our data support a model where the location of endocytosis serves as a spatial mark for septin structure assembly and that desensitization of the Gα delays its endocytosis sufficiently that septins are placed peripheral to the site of Cdc42 polarity.

## Introduction

Septins are evolutionarily conserved cytoskeletal filaments that are known for their roles as diffusion barriers, cortical actin regulators, and scaffolds that fortify membrane shape (Shuman and Momany, 2021; Spiliotis and McMurray, 2020). Septins are most well known for their role in mitosis, playing a part in the contractile ring and division plane control (Bridges and Gladfelter, 2015). Septins have nonmitotic roles as well, forming barriers and promoting curvature, functions important to morphogenesis of both the axon initial segment and the dendritic spine of neurons (Ewers et al., 2014; Hamdan et al., 2020; Xie et al., 2007). Septin structures form at the base of cilia and compartmentalize endothelial junctions (Kim et al., 2023; Palander et al., 2017). However, the molecular mechanisms by which cells organize the placement of nonmitotic septins are not well understood.

Much of our understanding of septins has come from the yeast Saccharomyces cerevisiae, where they were first discovered forming a double-ring diffusion barrier between mother and daughter yeast during mitosis. Outside of mitosis, septins are required for yeast mating projection morphogenesis (Bridges and Gladfelter, 2015; McMurray et al., 2011). At a high dose of pheromone, as when close to a mate, yeast form a tight mating projection called a shmoo that is bounded by septins at the base (Giot and Konopka, 1997; Longtine et al., 1998). At an intermediate dose of pheromone, as when farther from a mate, yeast undergo elongated growth towards the source of pheromone in a gradient tracking process that requires septins (Kelley et al., 2015; Segall, 1993).

The yeast mating response is driven by a G-protein coupled receptor (GPCR) signaling pathway that initiates transcription of mating genes, polarization of the actin cytoskeleton, and growth of the mating projection. The α-Factor pheromone is detected by the GPCR Ste2, which initiates signaling through the heterotrimeric G-protein comprised of the α (Gpa1), β (Ste4), and γ (Ste18) subunits (Alvaro and Thorner, 2016). While Gβγ signaling has traditionally taken center stage for regulating Cdc42-mediated polarity and MAPK signaling, Gα also controls downstream signaling. Active Gpa1 traffics to the endosome through its unique Ubiquitination Domain (UD), a 109 amino acid insertion not found outside of fungi (Dixit et al., 2014a). At the endosome, Gpa1 interacts with the PI3 Kinase Vps34, promoting PI(3)P production (Dixit et al., 2014a; Slessareva and Dohlman, 2006). We previously found that Gpa1 activity influences the spatial distribution of septins during the pheromone response and that septins are required for gradient tracking (Kelley et al., 2015).

Septin deposition on the membrane is controlled by the interplay Cdc42 effectors, including the septin chaperones called GTPase interactive components (Gic1 and Gic2), and the Cdc42 GTPase Activating Proteins (GAPs, Rga1, Rga2, and Bem3) (Khan et al., 2015). The Cdc42 GAPs bind the epsins Ent1 and Ent2, endocytic adaptor proteins that have been reported to lead to septin defects when mutated (Aguilar et al., 2006; Mukherjee et al., 2009).

During mitosis, it is thought that septins form a ring through the interplay of exocytosis and negative regulation by the CDC42 GAPs (Okada et al., 2013). Exocytic events push the membrane-associated septins away from the polarity site, helping to generate the well-known ring structure that septins form between mother and daughter yeast. The mechanism that controls the spatial distribution of septins during the pheromone response has not yet been elucidated but is influenced by the activity of Gpa1 (Kelley et al 2015). Gpa1 is desensitized through the actions of the regulator of G-protein signaling (RGS), Sst2, which serves as a GAP for Gpa1 (Apanovitch et al., 1998; Dohlman et al., 1996). Either deletion of the RGS or mutation of Gpa1 to disrupt RGS binding (*gpa1^G302S^*) leads to septin mislocalization to the site of polarity (DiBello et al., 1998; Kelley et al., 2015). *sst2^Q304N^* mutation leads to equivalent signaling through the Gβγ pathway as the *gpa1^G302S^* mutation but does not lead to the defects in septin organization (Dixit et al., 2014b; Kelley et al., 2015). Thus, the septin defect is mediated by Gpa1.

In this study, we aimed to understand how Gpa1 controls septin organization during the pheromone response. We hypothesized that Gpa1 influences septin organization through known septin regulators, including Gic1/2, Cdc42 GAPs, and epsins. To test this, we used gene deletions combined with microfluidics and live-cell fluorescence microscopy to examine the rescue of peripheral septin distribution in cells expressing the hyperactive Gα mutant *gpa1^G302S^*. Our findings revealed that an Ent1/2-Bem3-Gic1 axis mediates Gpa1’s control of septin distribution during the pheromone response. The involvement of epsins led us to hypothesize that the location of endocytosis may regulate septin organization. Supporting this, we developed a mathematical model showing that endocytic events could generate the observed septin patterns. We then experimentally disrupted clathrin-mediated endocytosis by deleting *END3* and found that endocytosis is necessary for Gpa1’s regulation of septins. Finally, we investigated the roles of two well-known pheromone-induced endocytic cargoes— the receptor Ste2 and Gpa1 itself. We discovered that endocytosis of either the receptor or Gpa1 can influence septin organization, but only the disruption of Gpa1 endocytosis can rescue the hyperactive *gpa1^G302S^* mutant.

## Methods

### Yeast Strains

The strains used in this study are listed in Table S1. All strains were constructed in the MATa haploid Saccharomyces cerevisiae parent strain, BY4741. All plasmids used in this study are included in Table S2. Polar cap (Bem1) and septin (Cdc3) proteins were tagged with GFP (Huh et al., 2003) and mCherry, respectively, at the native chromosomal locus through oligonucleotide-directed homologous recombination using primers listed in Table S3.

Cells were grown in rich medium (YPD) or Synthetic Complete +Dextrose medium (SCD) at 30°C unless otherwise indicated. PCR products were transformed into parent yeast strains using standard lithium acetate transformation (Burke et al., 2000). Individual colonies were isolated by growth on standard selective media (SC leu-, SC ura-, SC his-, or YPD G418+). Transformants were verified using fluorescence microscopy, sequencing, and/or PCR.

### Cloning

*gpa1*^Δ*UD-G302S*^ *Plasmid Cloning* The gpa1^ΔUD-G302S^ plasmid was constructed using the previously published pRS406-GPA1_truncΔUD (Dixit et al., 2014a), which we received as a gift from the Dohlman Lab. We mutagenized the pRS406-GPA1_truncΔUD using the site-directed mutagenesis kit (NEB) in combination with the primers AHM-14 and AHM-15 listed in table 3. The plasmid was sequenced via PlasmidSaurus (https://www.plasmidsaurus.com/) to confirm successful mutagenesis of the glycine at position 302 to serine. Following sequence confirmation, we linearized the plasmid for genomic integration using AflII (NEB), a silent mutation inserted previously. For cloning of the Gal-inducible Ste5-CTM plasmid, the galactose-inducible STE5 with the C-terminal membrane domain (CTM) of SNC2 was excised from the parent plasmid pGS5-CTM using *SacI-HF* and *ApaI* endonucleases (Pryciak and Huntress, 1998). pRSII416 plasmid (Chee and Haase, 2012) was linearized using the same enzymes, followed by a treatment of Antarctic phosphatase (New England Biolabs) to produce a new parent vector for the Gal-inducible Ste5-CTM fragment. Verification of the fragment and vector was performed using gel electrophoresis with a 0.8% TBE-agarose gel to confirm the size of the two products. The fragment and vector were then ligated together overnight at 16 °C using T4 DNA ligase. The ligation mixture was transformed into DH5α competent *E. coli* cells (NEB #C2987), and colonies were isolated by growth on LB agar plates supplemented with 0.1 mg/mL carbenicillin. Plasmids were extracted from these transformants and verified by restriction digest with *SacI-HF* and *ApaI*, followed by gel electrophoresis, and ultimately sequencing by PlasmidSaurus.

**Table 1:**
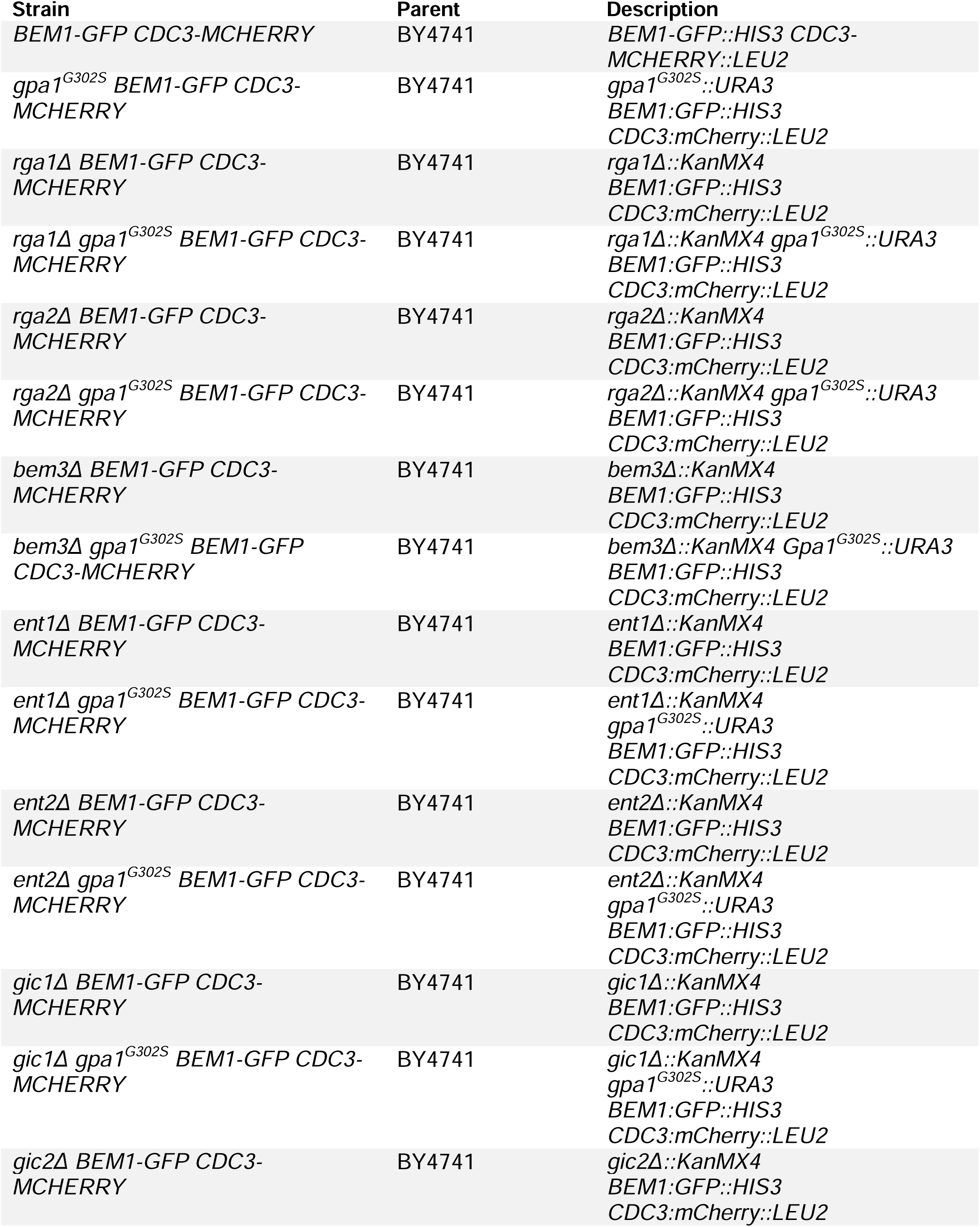
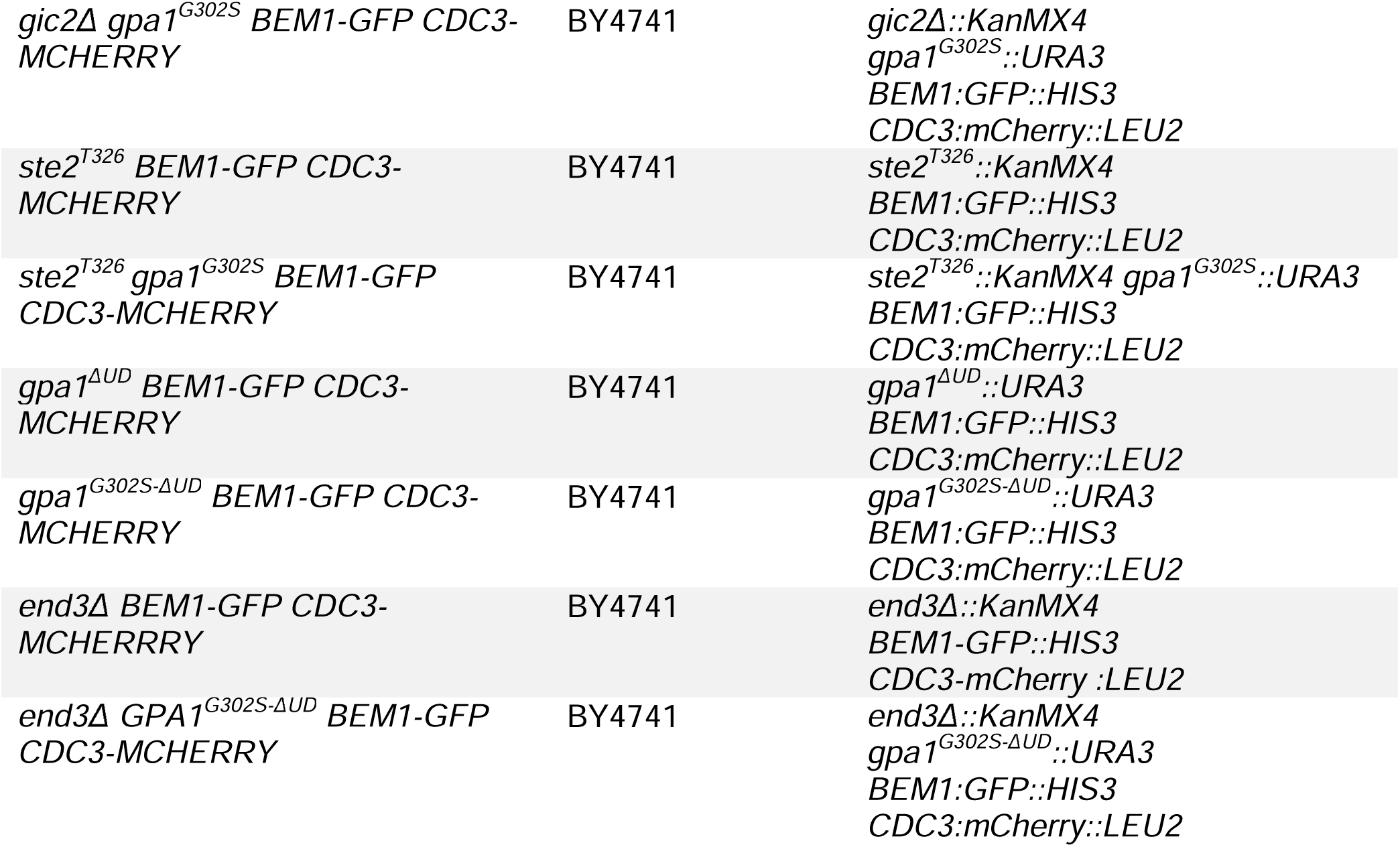
Strain List.

| Strain | Parent | Description |
| --- | --- | --- |
| *BEM1-GFP CDC3-MCHERRY* | BY4741 | *BEM1-GFP::HIS3 CDC3-MCHERRY::LEU2* |
| *gpa1<sup>G302S</sup> BEM1-GFP CDC3-MCHERRY* | BY4741 | *gpa1<sup>G302S</sup>::URA3*<br>*BEM1-GFP::HIS3*<br>*CDC3:mCherry::LEU2* |
| *rga1Δ BEM1-GFP CDC3-MCHERRY* | BY4741 | *rga1Δ::KanMX4*<br>*BEM1-GFP::HIS3*<br>*CDC3:mCherry::LEU2* |
| *rga1Δ gpa1<sup>G302S</sup> BEM1-GFP CDC3-MCHERRY* | BY4741 | *rga1Δ::KanMX4 gpa1<sup>G302S</sup>::URA3*<br>*BEM1-GFP::HIS3*<br>*CDC3:mCherry::LEU2* |
| *rga2Δ BEM1-GFP CDC3-MCHERRY* | BY4741 | *rga2Δ::KanMX4*<br>*BEM1-GFP::HIS3*<br>*CDC3:mCherry::LEU2* |
| *rga2Δ gpa1<sup>G302S</sup> BEM1-GFP CDC3-MCHERRY* | BY4741 | *rga2Δ::KanMX4 gpa1<sup>G302S</sup>::URA3*<br>*BEM1-GFP::HIS3*<br>*CDC3:mCherry::LEU2* |
| *bem3Δ BEM1-GFP CDC3-MCHERRY* | BY4741 | *bem3Δ::KanMX4*<br>*BEM1-GFP::HIS3*<br>*CDC3:mCherry::LEU2* |
| *bem3Δ gpa1<sup>G302S</sup> BEM1-GFP CDC3-MCHERRY* | BY4741 | *bem3Δ::KanMX4 Gpa1<sup>G302S</sup>::URA3*<br>*BEM1-GFP::HIS3*<br>*CDC3:mCherry::LEU2* |
| *ent1Δ BEM1-GFP CDC3-MCHERRY* | BY4741 | *ent1Δ::KanMX4*<br>*BEM1-GFP::HIS3*<br>*CDC3:mCherry::LEU2* |
| *ent1Δ gpa1<sup>G302S</sup> BEM1-GFP CDC3-MCHERRY* | BY4741 | *ent1Δ::KanMX4*<br>*gpa1<sup>G302S</sup>::URA3*<br>*BEM1-GFP::HIS3*<br>*CDC3:mCherry::LEU2* |
| *ent2Δ BEM1-GFP CDC3-MCHERRY* | BY4741 | *ent2Δ::KanMX4*<br>*BEM1-GFP::HIS3*<br>*CDC3:mCherry::LEU2* |
| *ent2Δ gpa1<sup>G302S</sup> BEM1-GFP CDC3-MCHERRY* | BY4741 | *ent2Δ::KanMX4*<br>*gpa1<sup>G302S</sup>::URA3*<br>*BEM1-GFP::HIS3*<br>*CDC3:mCherry::LEU2* |
| *gic1Δ BEM1-GFP CDC3-MCHERRY* | BY4741 | *gic1Δ::KanMX4*<br>*BEM1-GFP::HIS3*<br>*CDC3:mCherry::LEU2* |
| *gic1Δ gpa1<sup>G302S</sup> BEM1-GFP CDC3-MCHERRY* | BY4741 | *gic1Δ::KanMX4*<br>*gpa1<sup>G302S</sup>::URA3*<br>*BEM1-GFP::HIS3*<br>*CDC3:mCherry::LEU2* |
| *gic2Δ BEM1-GFP CDC3-MCHERRY* | BY4741 | *gic2Δ::KanMX4*<br>*BEM1-GFP::HIS3*<br>*CDC3:mCherry::LEU2* |
| *gic2Δ gpa1<sup>G302S</sup> BEM1-GFP CDC3-MCHERRY* | BY4741 | *gic2Δ::KanMX4*<br>*gpa1<sup>G302S</sup>::URA3*<br>*BEM1:GFP::HIS3*<br>*CDC3:mCherry::LEU2* |
| *ste2<sup>T326</sup> BEM1-GFP CDC3-MCHERRY* | BY4741 | *ste2<sup>T326</sup>::KanMX4*<br>*BEM1:GFP::HIS3*<br>*CDC3:mCherry::LEU2* |
| *ste2<sup>T326</sup> gpa1<sup>G302S</sup> BEM1-GFP CDC3-MCHERRY* | BY4741 | *ste2<sup>T326</sup>::KanMX4 gpa1<sup>G302S</sup>::URA3*<br>*BEM1:GFP::HIS3*<br>*CDC3:mCherry::LEU2* |
| *gpa1<sup>ΔUD</sup> BEM1-GFP CDC3-MCHERRY* | BY4741 | *gpa1<sup>ΔUD</sup>::URA3*<br>*BEM1:GFP::HIS3*<br>*CDC3:mCherry::LEU2* |
| *gpa1<sup>G302S-ΔUD</sup> BEM1-GFP CDC3-MCHERRY* | BY4741 | *gpa1<sup>G302S-ΔUD</sup>::URA3*<br>*BEM1:GFP::HIS3*<br>*CDC3:mCherry::LEU2* |
| *end3Δ BEM1-GFP CDC3-MCHERRY* | BY4741 | *end3Δ::KanMX4*<br>*BEM1-GFP::HIS3*<br>*CDC3-mCherry :LEU2* |
| *end3Δ GPA1<sup>G302S-ΔUD</sup> BEM1-GFP CDC3-MCHERRY* | BY4741 | *end3Δ::KanMX4*<br>*gpa1<sup>G302S-ΔUD</sup>::URA3*<br>*BEM1:GFP::HIS3*<br>*CDC3:mCherry::LEU2* |

**Table 2:** Plasmid List.

| **Plasmid Name** | **Vector Backbone** | **Description** |
| --- | --- | --- |
| **pRSII406-GPA1<sup>G302S</sup>** | pRSII406 | Integrating gpa1<sup>G302S</sup>::URA3 vector |
| **pRSII406-GPA1<sup>G302S-ΔUD</sup>** | pRSII406 | Integrating gpa1<sup>G302S-ΔUD</sup>::URA3 vector |
| **pRSII416-GAL-Ste5CTM** | pRSII416 | Galactose-inducible Ste5-CTM |
| **pFA6a-yo-linkEGFP-KanMX** | pFA6a | Addgene #44900 (Lee et al., 2013) |

**Table 3:**
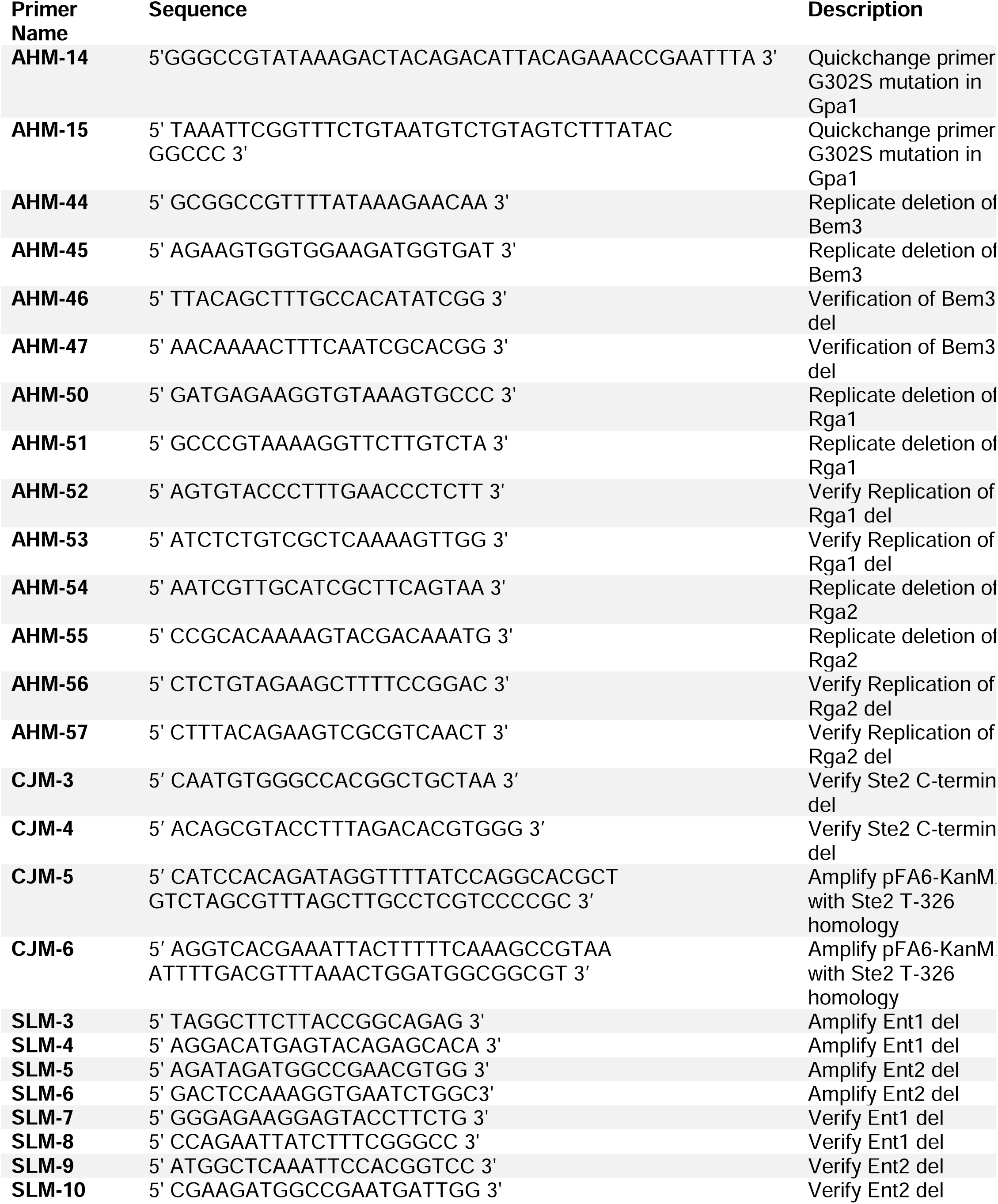
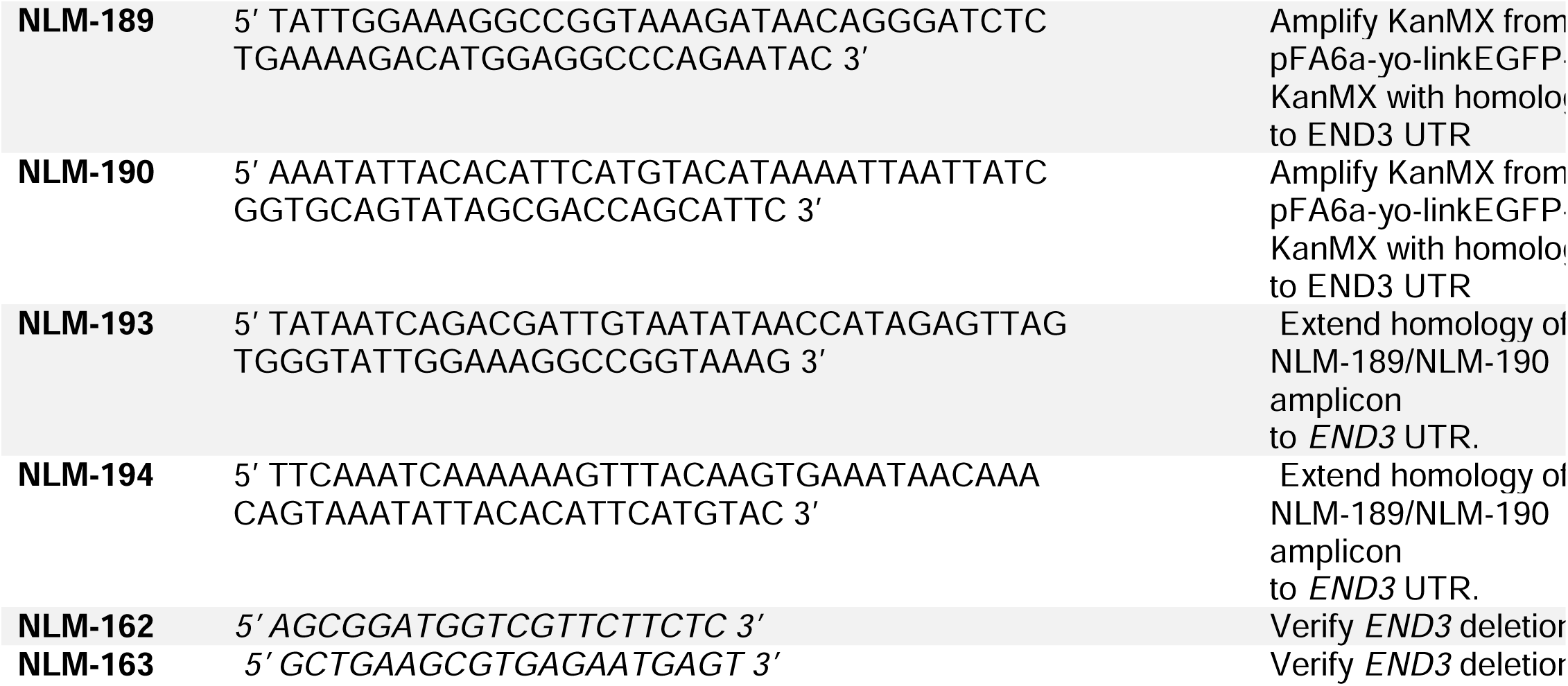
Primer List.

### Microfluidics Experiments

Microfluidic devices were made by mixing SYLGARD 184 silicone polymer at 10(part A):1(part B) and poured directly onto a microfluidics device mold (Suzuki et al., 2021) that was fabricated by the University of Maine Frontier Institute for Research in Sensor Technologies (FIRST, https://umaine.edu/first/). Microfluidic chambers were made as previously described (Suzuki et al., 2021) Molds and coverslips were exposed to oxygen plasma for 50 seconds in a Harrick Plasma Cleaner PDC32G (Harrick Plasma) followed by fusion of the device mold to the coverslip. After fusing, the device was then baked in an oven at 80°C for at least 20 minutes to facilitate fusion of the mold to the coverslip.

Cultures were grown in 0.45µm-filtered SCD to an OD600 between 0.1-0.8 at 30°C. Live-cell microfluidics experiments were performed using an IX83 (Olympus, Waltham MA) microscope and a Prime 95B CMOS Camera (Photometrics) controlled by Cell Sens 1.17 (Olympus). Fluorescence and Differential Interference Contrast (DIC) images were acquired using an Olympus-APON-60X-TIRF objective. Z-stacks of GFP and mCherry images were acquired using an X-Cite 120 LEDBoost (Excelitas). Cells were imaged in a “Dial-a-wave” based microfluidic device that allowed for rapid switching of media while yeast remained in place (Bennet et al., 2011; Dixit et al., 2014b; Suzuki et al., 2021). Pheromone-containing media contains AlexaFluor 647 dye at 1:8000 dilution (Life Technologies) and imaged in a single plane to verify proper flow within the chamber. Cells were imaged at 20-minute intervals for 12 hours in SCD with 300nM pheromone present in the media. Imaging settings were determined based on experimental needs and were replicated for repeat experiments.

### Ste5-CTM Experiments

Yeast cultures containing the Ste5-CTM (pRSII416) plasmid were diluted and grown overnight at 30 °C shaking in SC ura-media supplemented with 2% raffinose such that they did not exceed a saturating concentration (OD600 = 0.8). These cultures were then diluted to OD600 = 0.2, treated with either 0.5% galactose or 0.5% galactose and 10µM pheromone, and allowed to incubate for 4 hours shaking at 30 °C prior to imaging. Fluorescence microscopy was used to image Bem1-GFP and Cdc3-mCherry.

### Image Analysis

Images were deconvolved using Huygens Software (Scientific Volume Imaging, Hilversum, Netherlands). For all fluorescent images, deconvolution was carried out with a theoretical point-spread function, 5 signal-to-noise ratio (GFP images) or 7 signal-to-noise ratio (mCherry images), maximum 500 iterations, and a quality change threshold of 0.01 using the classical algorithm. All other settings were set to default. Images were saved as 16-bit TIFF with linked-scale. Cell masks were made alternatively through the use of Cellpose (www.cellpose.org) (Pachitariu and Stringer, 2022; Stringer et al., 2021) or by using a merged 8-bit RGB version of GFP and mCherry channels in FIJI. Septin localization analysis in MATLAB. All septin localization analysis was conducted using *whole_cell_cap_v4.m* for cell tracking and *combokeeper_v3.m* for line scan analysis Septin data was aligned to the polar cap profile using the *regalignraw.m* script (GitHub) and subsequently normalized as previously described (Kelley et al., 2015; Simke et al., 2022). In short, the minimum value in each profile was subtracted, and then the data was normalized to sum to 1. and 95% confidence intervals of the mean intensity were determined by bootstrapping. The referenced MATLAB scripts for these analyses are available on Github (https://github.com/Kelley-Lab-Computational-Biology/Spatial-Normalization).

### Classification of Septins

Symmetry was removed from the centered and normalized septin profiles by flipping any profiles with higher signal to the left of the center (location of peak Bem1) such that all profiles have higher signal to the right of the center using the MATLAB function *flipud()*. The complete asymmetric septin data set was loaded into a single variable and principal component analysis was performed with the MATLAB function *pca()*. We examined the explained variance, and found that the top 12 scores accounted for >95% of the variance, and we proceeded with data represented by the first 12 PCA scores. We then performed iterative K-means clustering with values of K ranging from 1 to 20 using the MATLAB function *kmeans()*. We compared the within-cluster sum of differences with increasing values of K – this tells us how much better the clustering works as the number of clusters increases. We then looked for a local maximum in the second derivative of this line, identifying K = 6 as the ideal number of clusters. For K > 6, improvements in the fit of the data to the clusters were lower.

### Agent-Based Model Description

The agent-based model of cargo and vesicle transport was made using MATLAB (Mathworks). The code is available on Github at https://github.com/Kelley-Lab-Computational-Biology/Trafficking-Model. The membrane consists of 100 bins, each 0.1 µm wide. The model has a time step of 0.01 seconds, and we ran the model for 200,000 steps per simulation.

-*Agents:* Individual cargo is tracked and can become phosphorylated with a probability of 5e-5 per time step (termed 1x phosphorylation). Phosphorylated cargo can be ubiquitinated, which happens with a probability of 0.01 per time step. Cargo which is phosphorylated and ubiquitinated is now competent for endocytosis. If it diffuses into an endocytic pit, it will be trapped and will no longer diffuse.

-*Diffusion:* The system is scaled such that diffusion can happen one bin at a time and recapitulate realistic receptor diffusion rates (Valdez-Taubas and Pelham, 2003). Movement probability incorporates cargo concentration in adjacent bins to weight diffusion probability to be more likely to move down the concentration gradient, as described previously (Azimi et al., 2011). The base probability to move one bin is 0.0024 per time step.

-*Vesicle Trafficking:* Exocytosis frequency has been measured by membrane capacitance at 2.4 events per minute in yeast that have been spheroplasted (Carrillo et al., 2015), and estimated to be 24 events per minute from microscopy data (Layton et al., 2011; Marco et al., 2007). Based on these two values, we chose an intermediate value of 8 events per minute, as yeast protoplasts may be experiencing cell wall stress that could decrease exocytosis. Endocytosis rate is set to twice the rate of exocytosis with a single 2 bin exocytic event and 2 independent single bin endocytic events. These rates and sizes are balanced to roughly maintain membrane size. Exocytic events push the surrounding bins away from the site of exocytosis, while endocytic events remove the affected bin, bringing the surrounding bins next to each other. Location of exocytic events is determined by the exocyst and Cdc42 (Brennwald, 2013), so we used a probability distribution based on the Bem1 and Exo84 distributions calculated during the pheromone response previously (Kelley et al., 2015). We used the probability distribution equal to [Bem1]^2^ x [Exo84] to account for the role of Cdc42-GTP (for which Bem1 serves as a marker) in both directing actin organization and promoting vesicle fusion. Location of endocytic events is determined by selecting the most mature endocytic pit. Whenever an endocytic pit is removed, a new one is formed at the site of the highest ubiquitinated cargo, consistent with the finding that the presence of cargo regulates endocytic pit location (Pedersen et al., 2020).

### Hierarchical Clustering and Similarity Analysis

We performed hierarchical clustering using the usage frequency of each septin class for each strain. Distances between datasets were calculated using the MATLAB *pdist()* function, and hierarchical clustering was performed with the *linkage()* function using the ‘average’ method. To assess the robustness of clustering, we performed bootstrapping to determine 95% confidence intervals for the clusters, following the method described by Mathworks (https://www.mathworks.com/help/bioinfo/ug/bootstrapping-phylogenetic-trees.html).

To examine the similarity in septin usage across strains, we projected the distance data onto a coordinate system where the x-axis represents the septin usage of wild-type (WT) and *gpa1^G302S^*strains. The distance between WT and *gpa1^G302S^* on this axis was maintained and centered around 0. Each strain was placed on the plot based on its calculated distance from both WT and *gpa1^G302S^*, using a method that identifies the intersection of two circles defined by these distances. Specifically, the distances served as radii from the two comparison strains, resulting in a single x-coordinate and both a positive and negative y-coordinate where the circles intersected. We recorded the positive y-coordinate for each strain. In this analysis, the x-axis represents Gpa1 activity, ranging from WT to hyperactive, while the y-axis captures orthogonal components of septin organization. We used bootstrapping to calculate 95% confidence intervals for the (x, y) positions of each strain.

### Halo Assays

Halo assays were performed as described previously (Hoffman et al., 2002). In short, α-factor pheromone was dissolved in water and the indicated amount was pipetted onto paper disks (0.1, 0.3, 1, and 3 µg). These disks were placed on a YPD agar plate containing a thin layer of yeast culture mixed with molten agar. Plates were incubated at 30°C for 24 hours and then photographed. The size of the zone of exclusion of growth for each strain was measured in FIJI.

### Use of AI

Some sections of text were edited with the assistance of ChatGPT 4o.

## Results

### Gpa1 regulation of septins occurs in addition to a default septin regulation pathway in response to pheromone

We hypothesized that Gpa1 control of septins would use a subset of known regulatory proteins. Our strategy to uncover this signaling pathway was to delete genes involved in septin regulation (Figure 1A) from cells expressing either WT Gpa1 or the hyperactive gpa1^G302S^ mutant and look for a rescue of the septin defects caused by the Gpa1 mutant. To investigate septin dynamics throughout the pheromone response, we employed live-cell microscopy in a microfluidic device. Cells were imaged in the presence of 300 nM pheromone for 12 hours in 20-minute intervals (Figure 1B, C). To quantify the distribution of septins along the periphery of the cell, we spatially normalized our septin profiles to the center of the polar cap – indicated by peak fluorescence of Bem1-GFP, as previously described (Kelley et al., 2015; Simke et al., 2022). We then averaged normalized septin distributions from all time points and bootstrapped 95% confidence intervals (indicated by shaded regions); non-overlapping confidence intervals indicate statistically significant differences for p < 0.05. This method for average septin organization relative to the polar cap is used throughout this study. We average all cells from different experiments together; experiment-to-experiment variability in averages for all experiments is shown in Supplemental Figure S1. Consistent with our previous work, septins form structures at the base of the mating projection in WT cells. However, hyperactive *gpa1^G302S^* mutants frequently show polar cap (Bem1) and septin (Cdc3) overlapping (Figure 1C). In quantitation, this leads to an increase in average septin concentration at the center of the polar cap as opposed to the well-separated peaks of septins ∼ 2 µm from the center of the polar cap in WT cells (Figure 1D). Average kymographs of Cdc3 and Bem1 for all experiments can be found in Supplemental Figures S2, S3, and S4.

**Figure 1.**
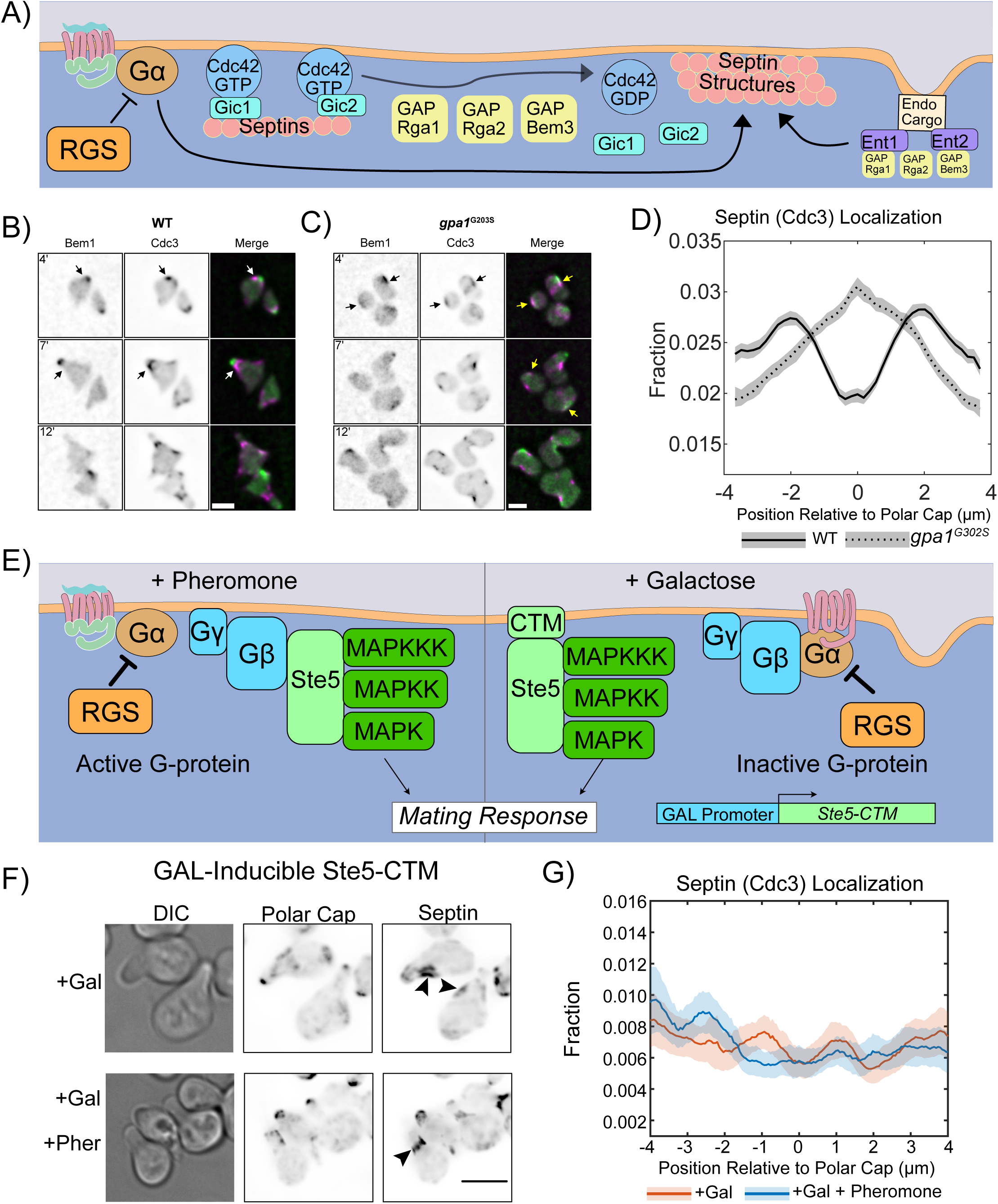
Pheromone-induced septin accumulation occurs at the center of the polar cap in a Hyperactive Gα mutant. A) Schematic of key players in septin deposition in yeast. Cdc42-GTP binds to GICs, which are required for septin recruitment. GAP activity promotes Cdc42 hydrolysis of GTP to GDP and septin structure formation. GAPs interact with the endocytic adapter proteins Ent1 and Ent2, which also impact septin organization through an unknown mechanism.B) Representative images of WT cells expressing endogenous Bem1-GFP (Polar Cap) and Cdc3-mCherry (Septin) exposed to 300 nM pheromone in a microfluidic chamber for 12hrs. Shown are images of cells at 4, 7, and 12 hours of pheromone treatment. White arrows indicate septins organized at the base of projections, away from the polar cap. Scale bar represents 5µm. C) Representative images of *gpa1^G302S^* yeast expressing endogenous Bem1-GFP (Polar Cap) and Cdc3-mCherry (Septin) exposed to saturating pheromone in a microfluidic chamber for 12hrs. Shown are images of cells at 4, 7, and 12 hours of pheromone treatment. Yellow arrows indicate areas of polar cap and septin colocalization. Scale bar represents 5µm. D) Time-averaged quantification of the fraction (y-axis) of septin and its distribution relative to the polar cap (x-axis). Lines are representations of time-averaged data (37 time points; 20min intervals for 12hrs) and shaded regions surrounding lines are 95% confidence intervals. Lines are representative of 4 combined experiments (WT n = 89 cells and *gpa1^G302S^* n = 98 cells). E) Diagram of how the galactose-induced Ste5-CTM construct functions to initiate the mating response without active Gpa1 or recruitment of endogenous Ste5 to the Gβγ. F) Representative images of cells expressing the GAL-inducible Ste5-CTM that activates the mating pathway without activation of Ste2, Gpa1, or Ste4/Ste18. Cells were treated with 0.5% galactose or 0.5% galactose & 10 µM pheromone for 4 hours. Arrows indicate sites of septin organization at the base of the mating projection. G) Quantitation of septin organization in cells expressing the GAL-inducible Ste5-CTM. Shaded area represents the 95% confidence intervals. n = 128 for +Gal and n = 139 for +Gal +Pheromone, across two separate experiments.

We previously found that hyperactive Gpa1 dysregulates septins during mating projection formation. However, we do not know if Gpa1 activation is required for normal septin organization. Septins are organized into a ring structure during mitosis in both haploids and diploids, despite low levels of Gpa1 in diploids (de Godoy et al., 2008), so we would anticipate that mechanisms for septin organization during cell division are independent of Gpa1. We tested whether Gpa1 activation is necessary for septin organization during the pheromone response by initiating MAPK signaling and Cdc42-mediated polarity using an inducible mutant MAPK scaffold, Ste5, modified on its C-terminus to contain a transmembrane domain (Ste5-CTM) (Pryciak and Huntress, 1998). Expression is controlled by a GAL promoter such that the pheromone pathway is inducible by galactose without activating the receptor Ste2 or activation of the hetero-trimeric G-protein, Gpa1(Gα), Ste4(Gβ), and Ste18(Gγ) (Figure 1E). We grew these cells in raffinose and then treated them with either 0.5% galactose for 4 hours or 0.5% galactose and 10 µM pheromone for 4 hours and imaged them. Cells were able to place septin structures at the base of the mating projection in the presence of galactose alone or with both galactose and pheromone (Figure 1F). When we examine the quantitation (Figure 1G), we find that Ste5-CTM expression generates two small peaks of septin intensity ∼1 µm from the center of the polar cap. When pheromone is added in addition to Ste5-CTM expression, localization is less well defined, with an asymmetric distribution and a peak further from the center of the polar cap. Although the septin structures formed are not as robust or consistent as those in a WT cell stimulated with pheromone, we can conclude that septin organization does not require active Gpa1. Thus, there is a baseline septin organization pathway that is Gpa1 independent.

### Quantitative Classification of Septin Organization in Mating Projections

The average septin profiles for WT and *gpa1^G302S^* presented in Figure 1D appears symmetric; however, the underlying data is asymmetric and heterogeneous as shown in the kymographs of individual cells for WT (Supplemental Figure S5) and *gpa1^G302S^* (Supplemental Figure S6). A mating yeast exhibits rotational symmetry around the axis of its mating projection. When we average data that is asymmetric to the right of the yeast with data that is asymmetric to the left, the result is a symmetric average, as shown in Figure 2A. To analyze the data in a way that more accurately reflects the underlying variations, we have developed a method for classifying the shape of the septin organization. In short, we remove the symmetry from the data set, reflecting profiles across the center of the polar cap such that the higher intensity side of the profile is always to the right (Figure 2B). We then perform a principal component analysis (PCA), followed by a K-means clustering with the first 12 scores of the PCA (accounting for >95% of the variance in the dataset). We found that 6 classes were ideal, as values of K larger than 6 led to diminishing improvements in the within-cluster sum of differences. We performed this for all septin profiles measured in this study (n = 72,222 distinct profiles across all strains and time points).

**Figure 2.**
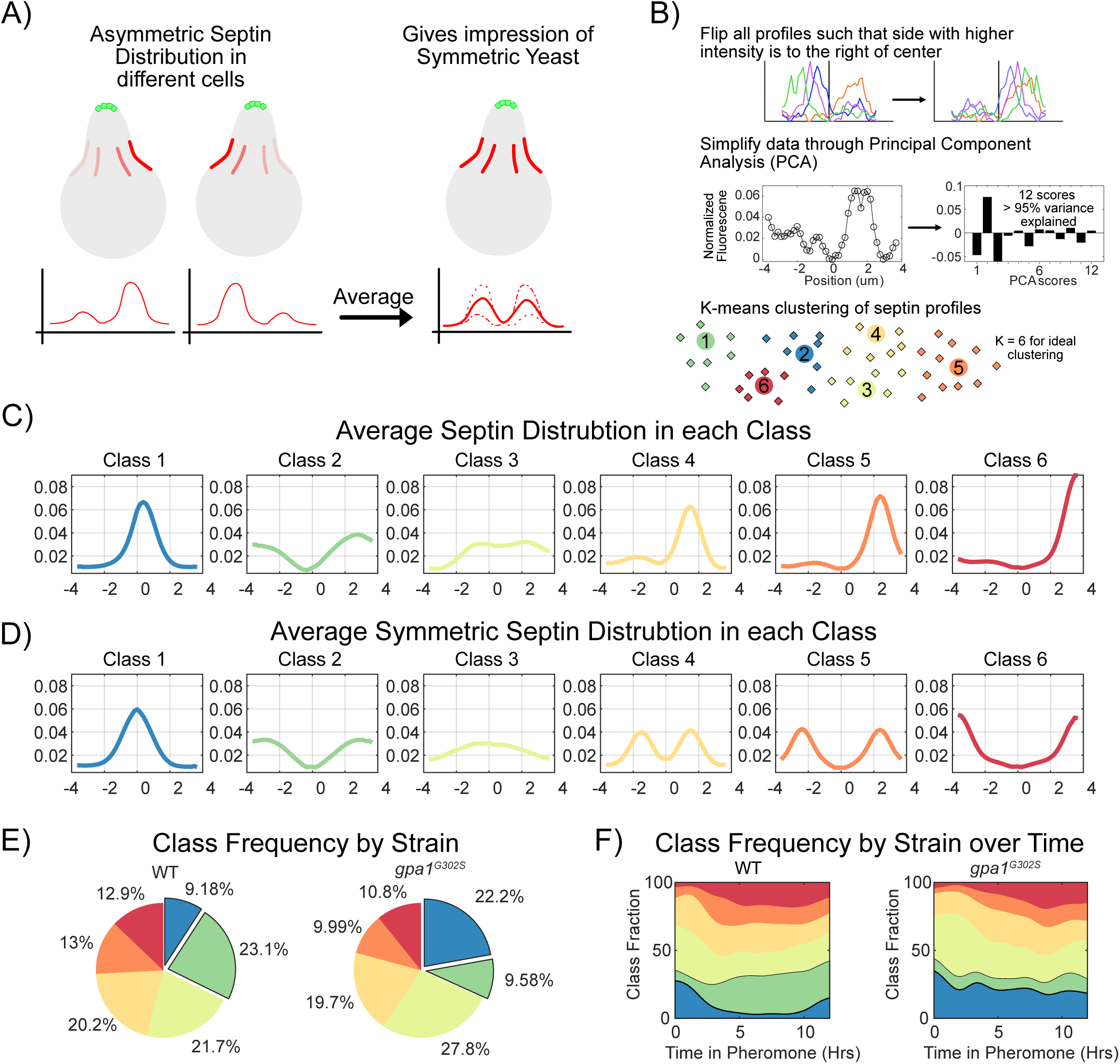
Classification of septin organization. A) The asymmetric distribution of septins in a cell with a rotational axis of symmetry means that septins on the right side of the cell and septins on the left side of the cell may have the same asymmetric localization. When data like this is averaged, it gives the appearance of symmetrical organization of septins. B) Symmetry was removed from the complete septin dataset (all data in this study) by horizontally flipping any septin profile that had higher intensity on the left of the polar cap, such that all septin profiles in the dataset are higher intensity to the right of the polar cap. All septin profiles were then simplified by principal component analysis and represented by the top 12 scores, which represented > 95% of the variance in the dataset. We then performed K-means clustering on the PCA-simplified dataset, identifying K=6 as the point where additional classes would yield diminishing returns. C) Graphs of the average of the members of each class using the asymmetric dataset. D) Graphs of the average of the members of each class using the symmetric dataset. E) Frequency of each septin class used in WT and *gpa1^G302S^*cells, based on the experiments from figure 1. Strikingly, the differences are mostly limited to class 1 and class 2 usage. F) Class usage over time for WT and *gpa1^G302S^* cells shows that class 1 is common in WT cells early in the pheromone response but transitions to use the more symmetric class 2 usage. By contrast, this transition into class 2 usage in *gpa1^G302S^* does not occur.

The averages of the members of each class are shown in Figure 2C, while Figure 2D shows what the average of each class is if the symmetry is not removed. Class 2 is the most symmetric of the classes, showing the expected peaks of septin intensity on both sides of the polar cap, what one might think of as the normal localization of septins during the pheromone response. Classes 4, 5, and 6 each show a strong localization to one side of the polar cap, with a smaller amount of septin signal on the other side. The averages of the symmetrical data make this look similar to Class 2, but in reality, these represent asymmetrical septin organization.

Having classified every septin profile, we can now consider the frequency at which septins within each class are used by different strains (Figure 2E). We find that WT and *gpa1^G302S^* strains show remarkable similarity in the use of Classes 3, 4, 5, and 6. Their differences arise from the switch between high class 2 and low class 1 usage in WT cells, to low class 2 and high class 1 usage in *gpa1^G302S^*. We can further look at how different classes are used over time (Figure 2F). At the beginning of the pheromone response, most cells are still going through mitosis, and class 1 and class 3 are the dominant classes being used. As WT cells progress through the mating response, they shift to more class 2 usage, creating symmetric septin structures around their polar cap. At later time points, classes 5 and 6 become more prevalent, indicating a transition to asymmetric septin organization and a polar cap that is getting further from the septin structures. In the *gpa1^G302S^*strain these trends are similar, except for the transition from class 1 into class 2 usage. Instead, gpa1^G302S^ cells keep a high use of Class 1, and their low use of Class 2 is consistent throughout the response. The frequency of septin class usage gives us a framework to quantitatively compare the similarities and differences between strains for this study.

### Gic1, but not Gic2, contributes to Gpa1-driven septin organization

The Cdc42-interacting paralogs, Gic1 and Gic2, are Cdc42-GTP binding proteins involved in the recruitment of septins to the polarity site during mitotic budding and in septin bundling activity *in vitro* (Figure 3A) (Iwase et al., 2006; Sadian et al., 2013). We constructed single-gene-deletions of Gic1 or Gic2 from WT and *gpa1^G302S^* strains and examined the polar cap (Bem1-GFP) and septin (Cdc3-mCherry) distribution (Figure 2B, C, D). In the presence of WT *GPA1*, neither deletion led to obvious morphological changes. In the presence of gpa1^G302S^, cells were rounded in both gic1Δ and gic2Δ. The average distribution of septins in gic1Δ has more signal peripheral to the polar cap, while in gic2Δ cells, the septin average is coincident with the center of the polar cap (Figure 3D).

**Figure 3.**
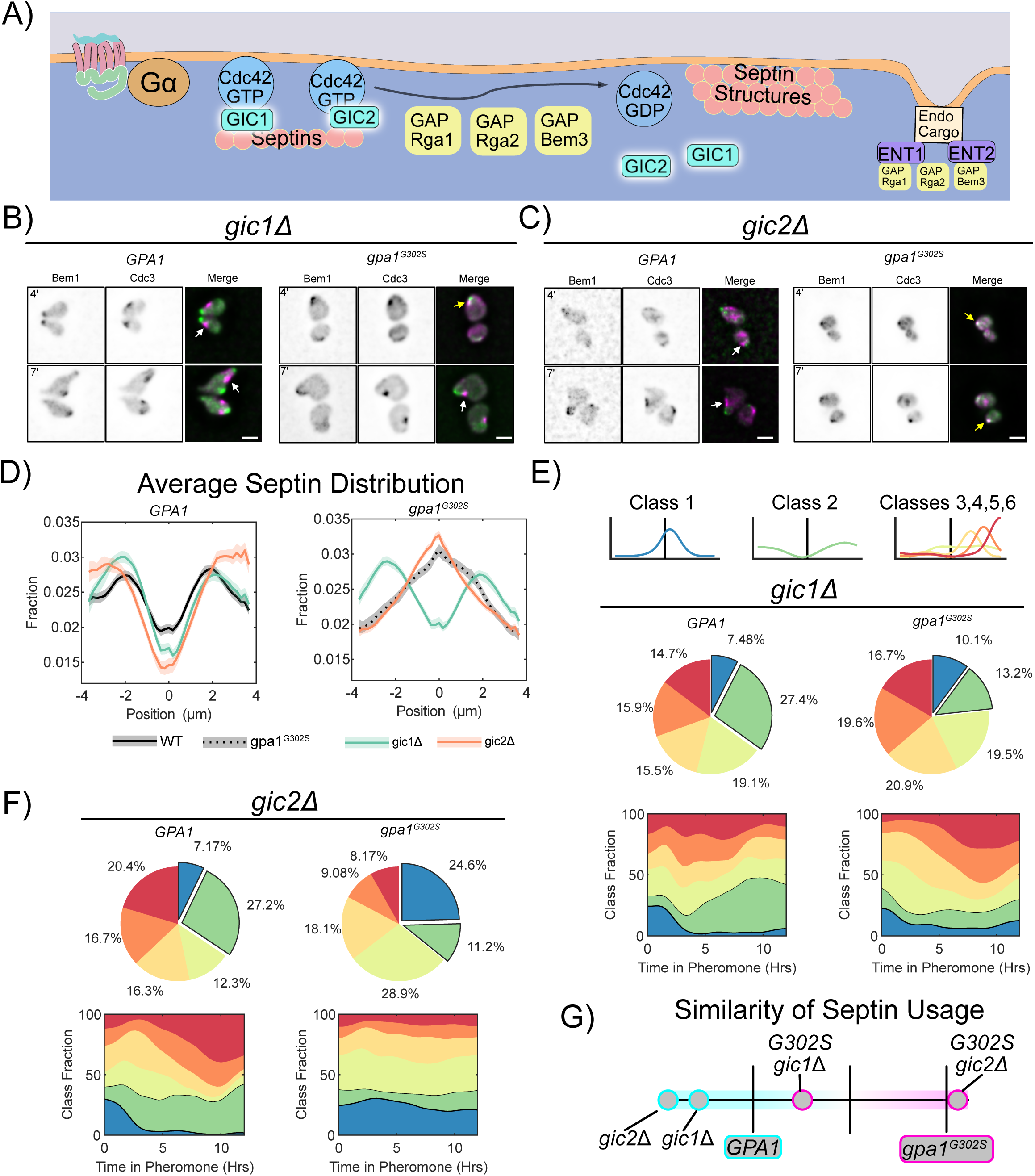
Gα control of septins is preferentially mediated by Gic1 rather than Gic2. A) Schematic showing Gic1 and Gic2 in septin organization. B) Representative images of *gic1*Δ *BEM1-GFP CDC3-mCherry* cells with WT *GPA1 or gpa1^G302S^* at 4 and 7 hours of treatment with 300nM pheromone. Scale bar represents 5µm. C) Representative images of *gic2*Δ *BEM1-GFP CDC3-mCherry* cells with WT *GPA1 or gpa1^G302S^* at 4 and 7 hours of treatment with 300nM pheromone. Scale bar represents 5µm. D) Average septin profiles from 12-hour microfluidic experiments, normalized to the center of the polar cap, for each of the indicated strains in *GPA1* or *gpa1^G302S^*backgrounds. WT *GPA1* and *gpa1^G302S^*without further gene deletions are from figure 1 and reproduced for ease of comparison. Shaded area represents 95% confidence intervals. Lines are representative of at least 2 combined experiments: *gic1*Δ n= 56 cells, *gpa1^g302s^ gic1*Δ n = 117 cells, gic2Δ n = 51 and *gpa1^G302S^ gic2*Δ n = 221 cells). E) Septin class usage in gic1Δ cells, with and without the *gpa1^G302S^* mutant, overall (pie charts) and over time (stacked graph). F) Septin class usage in gic2Δ cells, with and without the *gpa1^G302S^* mutant, overall (pie charts) and over time (stacked graph). G) Similarity of the septin usage for each deletion strain compared to WT *GPA1* or *gpa1^G302S^* alone based on septin class frequency distance calculation.

Interestingly, both Gic deletions led to decreased usage of Class 1 septin distribution and increased usage of Class 2 septin distributions in the WT *GPA1* background. In the presence of *gpa1^G302S^*, however, deletion of Gic1 leads to a drastic reduction in Class 1 septins, and an increase in Classes 5 and 6, while deletion of Gic2 has minimal impact on septin organization (Figure 3 E, F). When we compare the similarity of the septin class usage among these strains (Figure 3G), we see that deletion of Gic1 from *gpa1^G302S^* cells leads to septin organization that is closer to WT *GPA1* than that of *gpa1^G302S^*. In contrast, deletion of Gic2 has minimal impact. Thus, Gic1 has a role in septin organization during the pheromone response distinct from that of Gic2 and is required for Gpa1 control of septins.

### The Cdc42 GAP, Bem3, mediates Gpa1 control of septin structures

Cdc42 GAP activity promotes septin structure assembly on the plasma membrane (Figure 4A) (Gladfelter et al., 2002; Khan et al., 2015). Yeast express three Cdc42 GAPs: Rga1, Rga2, and Bem3 (Smith et al., 2002). Deletion of Rga1, Rga2, or Bem3 from WT GPA1 cells had minimal effects on cell morphology and average septin localization (Figure 4 B, C, D, E). In the *gpa1^G302S^* background, deletion of *RGA1* showed no discernable effect on morphology and only a mild impact on average septin localization. Deletion of *RGA2* resulted in fewer rounded cells and an average septin localization falling between WT and *gpa1^G302S^*. Deletion of *BEM3* improved morphology relative to *gpa1^G302S^*, though cells remained abnormal. Notably, the average septin profile in bem3Δ exhibited a more WT-like distribution.

**Figure 4.**
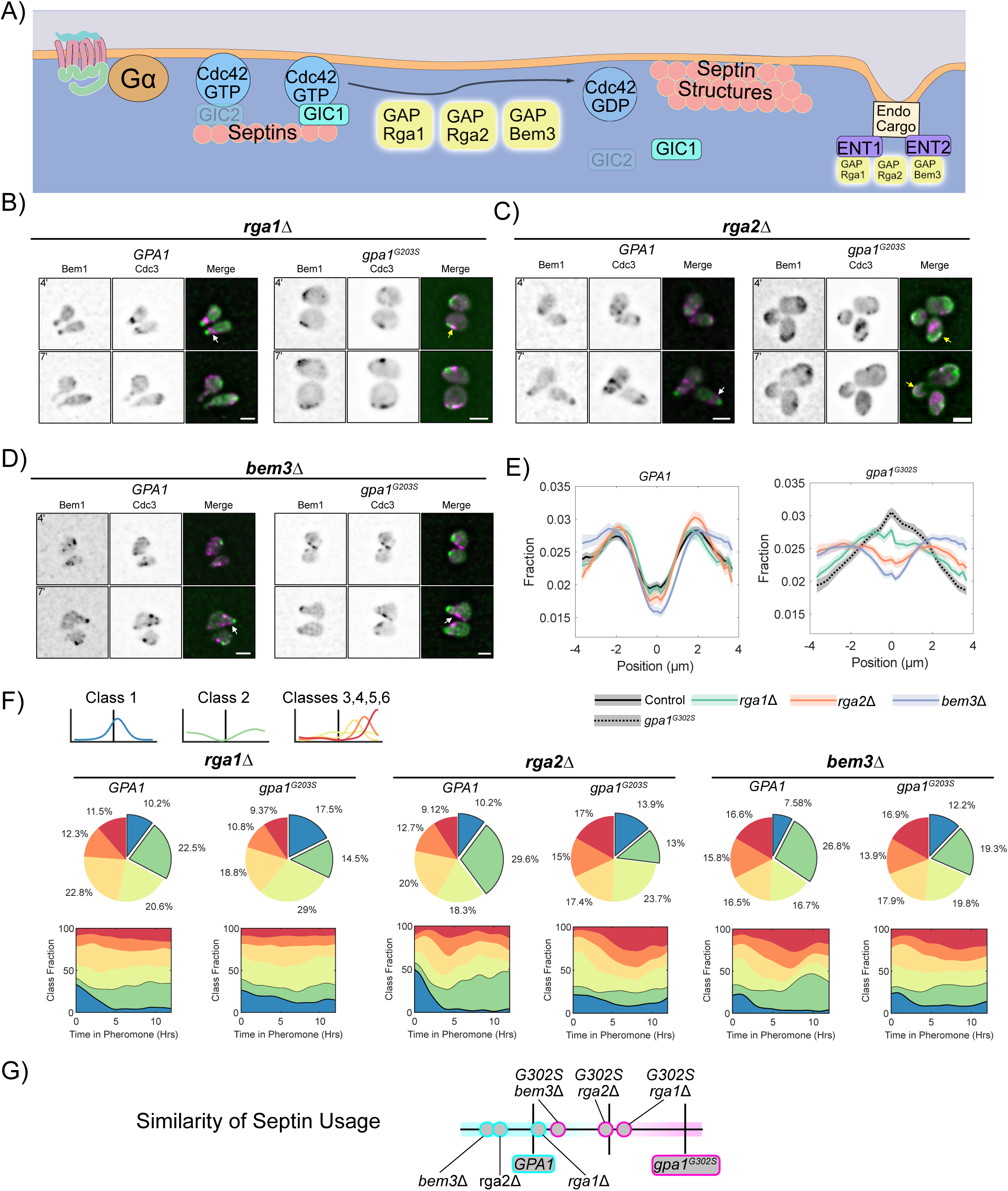
Gα control of septins is preferentially mediated by the Cdc42GAP Bem3, rather than Rga1 or Rga2. A) Schematic showing the Cdc42 GAPs Rga1, Rga2 and Bem3 in septin organization. B) Representative images of *rga1*Δ *BEM1-GFP CDC3-mCherry* cells with WT *GPA1 or gpa1^G302S^* at 4 and 7 hours of treatment with 300nM pheromone. Scale bar represents 5µm. C) Representative images of *rga2*Δ *BEM1-GFP CDC3-mCherry* cells with WT *GPA1 or gpa1^G302S^*at 4 and 7 hours of treatment with 300nM pheromone. Scale bar represents 5µm. D) Representative images of *bem3*Δ *BEM1-GFP CDC3-mCherry* cells with WT *GPA1 or gpa1^G302S^* at 4 and 7 hours of treatment with 300nM pheromone. Scale bar represents 5µm. E) Average septin profiles from 12-hour microfluidic experiments, normalized to the center of the polar cap, for each of the indicated strains in *GPA1* or *gpa1^G302S^*backgrounds. WT *GPA1* and *gpa1^G302S^*without further gene deletions are from figure 1 and reproduced for ease of comparison. Shaded area represents 95% confidence intervals. Lines are representative of at least 2 combined experiments: *rga1*Δ n= 55 cells, *gpa1^g302s^ rga1*Δ n = 102 cells, *rga2*Δ n = 51 and *gpa1^G302S^ rga2*Δ n = 103 cells, bem3Δ n = 73, *gpa1^G302S^ bem3*Δ n = 122. F) Septin class usage in *rga1*Δ, *rga2*Δ, and *bem3*Δ cells, with and without the *gpa1^G302S^* mutant, overall (pie charts) and over time (stacked graph). G) Similarity of the septin usage for each deletion strain compared to WT *GPA1* or *gpa1^G302S^* alone based on septin class frequency distance calculation.

Analysis of class usage in *GPA1* cells revealed that deletion of *RGA1* had minimal impact, whereas deletion of *RGA2* or *BEM3* slightly increased usage of Class 2 septin profiles, particularly at later time points (Figure 4F). In *gpa1^G302S^* cells, deletion of RGA1 did little to stop aberrant Class 1 septin usage. In contrast, deletion of *RGA2* reduced Class 1 but did not promote a corresponding increase in Class 2. Only deletion of *BEM3* significantly reduced Class 1 usage while increasing Class 2 usage, resulting in septin usage more like WT.

When comparing the similarity of septin class usage among these strains (Figure 4G), *BEM3* deletion produced the strongest rescue, while *RGA2* deletion provided partial rescue, with results falling almost exactly between WT and gpa1^G302S^ septin usage. These results suggest that Rga1 plays little role in septin organization during the pheromone response, Rga2 has a minor role, and Bem3 is a key contributor to Gpa1-mediated control of septins.

### The endocytosis adaptors Ent1 and Ent2 direct Gpa1 dependent pheromone-induced septin organization

Cdc42 GAPs bind directly to the epsin endocytic-adaptor proteins, Ent1 and Ent2, in an interaction thought to couple Cdc42 regulation to endocytosis (Aguilar et al., 2006). Additionally, epsins are known to differentially contribute to septin organization during cell division (Mukherjee et al., 2009). We hypothesized that epsins may also contribute to septin organization during the pheromone response (Figure 5A). As above, we investigated septin organization in WT *GPA1* and *gpa1^G302S^* mutants lacking either Ent1 or Ent2 via live cell imaging (Figure 5B, C). Notably, both *ent1*Δ and *ent2*Δ frequently made bean-shaped cells with asymmetric septin organization to one side of the polar cap. Average septin organization appears peripheral to the polar cap, indicating that both strains stop gpa1^G302S^ from placing septins close to the polar cap. Septin class usage was fairly similar between *gpa1^G302S^ ent1*Δ and *gpa1^G302S^ ent2*Δ, but the latter showed a larger increase in Class 2 septin usage at later time points, mimicking WT septin usage more closely (Figure 5E, F). This bears out in the analysis of the similarity of the septin usage, with g*pa1^G302S^ ent2*Δ being closer to WT than *gpa1^G302S^ ent1*Δ (Figure 5G). This is notable because ent2 preferentially interacts with bem3 (Costakes et al., 2013; Mukherjee et al., 2009). Thus, while both Ent1 and Ent2 are mediators of Gpa1 control of septin distribution during the pheromone response, Ent2 appears to play a larger role.

**Figure 5.**
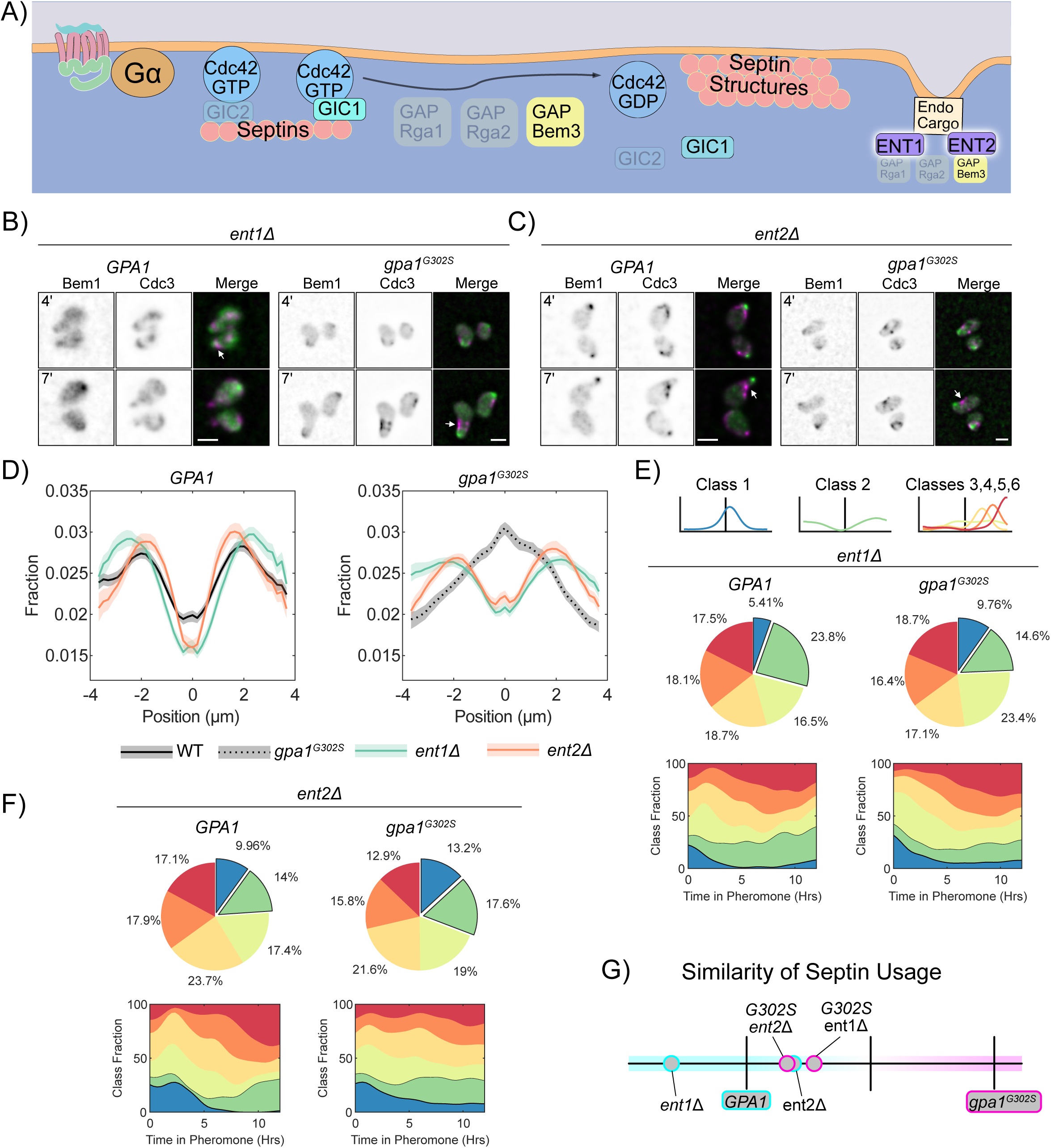
Gα control of septins can be mediated by either of the endocytic adaptor proteins, Ent1 or Ent2. A) Schematic showing Ent1 and Ent2 in septin organization. B) Representative images of *ent1*Δ *BEM1-GFP CDC3-mCherry* cells with WT *GPA1 or gpa1^G302S^* at 4 and 7 hours of treatment with 300nM pheromone. Scale bar represents 5µm. C) Representative images of *ent2*Δ *BEM1-GFP CDC3-mCherry* cells with WT *GPA1 or gpa1^G302S^*at 4 and 7 hours of treatment with 300nM pheromone. Scale bar represents 5µm. D) Average septin profiles from 12-hour microfluidic experiments, normalized to the center of the polar cap, for each of the indicated strains in *GPA1* or *gpa1^G302S^*backgrounds. WT *GPA1* and *gpa1^G302S^*without further gene deletions are from figure 1 and reproduced for ease of comparison. Shaded area represents 95% confidence intervals. Lines are representative of at least 2 combined experiments: *ent1*Δ n= 53 cells, *gpa1^g302s^ ent1*Δ n = 118 cells, *ent2*Δ n = 50 cells and *gpa1^G302S^ ent2*Δ n = 162 cells). E) Septin class usage in ent1Δ cells, with and without the *gpa1^G302S^*mutant, overall (pie charts) and over time (stacked graph). F) Septin class usage in ent2Δ cells, with and without the *gpa1^G302S^* mutant, overall (pie charts) and over time (stacked graph). G) Similarity of the septin usage for each deletion strain compared to WT *GPA1* or *gpa1^G302S^* alone based on septin class frequency distance calculation.

### An agent-based model of vesicle trafficking shows that location of endocytosis could explain septin organization

Strong rescue by both endocytic adaptor proteins Ent1 and Ent2 led us to hypothesize that endocytosis itself may be able to regulate the location of septin organization. As the rescue of septin organization in each experiment has been through a loss of function (deletion) mutation, we would presume that our septin rescues are due to a slowing of a process. Thus, the gpa1^G302S^ defects must be driven by speeding up a process. If septin organization is linked to endocytosis, then increases in the rate at which cargo becomes competent for endocytosis could change the location of endocytic events and give rise to the septin redistribution to the polar cap we see in mutant cells. To test that rates of endocytic competency could drive these patterns, we developed an agent-based stochastic model of protein trafficking on the membrane. This model is described in Figure 6. Cargo is delivered to the membrane, diffuses, undergoes endocytosis, and is tracked spatially over time. Like the receptor, Ste2, it has a probability to become phosphorylated, and phosphorylated cargo has a probability to become ubiquitinated, making it competent for endocytosis (Figure 6A) (Lu et al., 2016). These reactions occur in a discretized 1-dimensional membrane that can grow and shrink with endo- and exocytosis (Figure 6B). The location of exocytosis is based upon the distributions of Bem1 and Exo84 that we have previously measured (Figure 6C) (Kelley et al., 2015). The number of endocytic pits is kept constant, such that when an endocytic pit is endocytosed, a new pit forms at a new site based upon the amount of ubiquitinated cargo, consistent with polarized endocytic cargo driving the polarization of endocytosis (Figure 6D) (Pedersen et al., 2020). Diffusion within the membrane is carried out as described by Azimi et al, where the direction of diffusion is biased by the occupancy of the adjacent bins such that diffusion into bins with fewer molecules is more likely (Figure 6E) (Azimi et al., 2011). Ubiquitinated cargo that enters an endocytic pit becomes trapped there, and eventually will be internalized (Figure 6F). We performed simulations where we varied the phosphorylation rate and examined the distribution of ubiquitinated cargo and the location of endocytic events.

**Figure 6.**
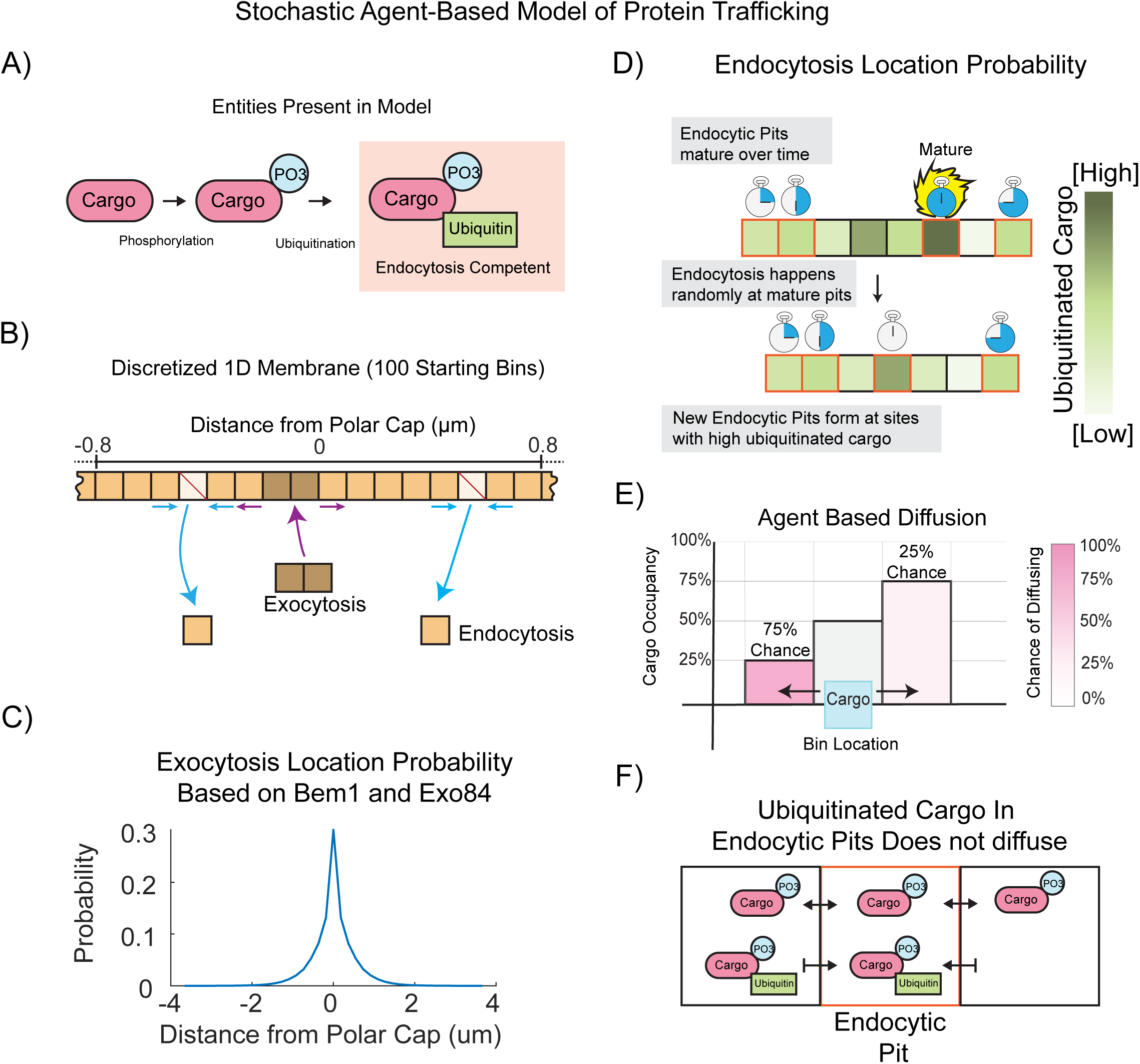
A stochastic agent-based model of protein trafficking. Schematic representation of the mathematical model of protein trafficking in which, (A) naïve cargo (unmodified) is first phosphorylated, then ubiquitinated. Upon ubiquitination, the cargo is considered “endocytically competent”. (B) The plasma membrane is represented by a 1-dimensional line segmented using 100 initial bins. Bins are removed via endocytosis and new bins are added via exocytosis. (C) The location of exocytosis is restricted to the center of the membrane with the given probability distribution. (D) In contrast, the location of endocytosis depends on two factors: location of endocytic pit formation and endocytic pit maturity. Endocytic pits form at sites with a high concentration of endocytosis competent cargo. Endocytic pits cannot diffuse, but their relative position on the membrane changes as bins are added and removed by vesicle trafficking. (E) To determine the directionality of cargo diffusion we employed an agent-based probability model. Briefly, the probability a cargo will diffuse to a bin is inversely correlated with how much cargo is currently in that bin. (F) Finally, once a ubiquitinated cargo enters an endocytic pit, it can no longer diffuse, leading to accumulation of endocytic cargo.

To investigate the effects of increased cargo endocytosis rates, we varied the rate of phosphorylation in our mathematical model and ran multiple simulations for each condition. Simulations tracked cargo location and modification on the cell surface over 30+ minutes of model time (Figure 7A). Increasing the phosphorylation rate from baseline (1x) to 5-fold or 25-fold accelerated the licensing of cargo for endocytosis (phosphorylation and ubiquitination), causing cargo to spend less time on the membrane before being internalized. As a result, the distribution of both total cargo (Figure 7A) and ubiquitinated cargo (Figure 7B) shifted closer to the center of the polarity site.

**Figure 7.**
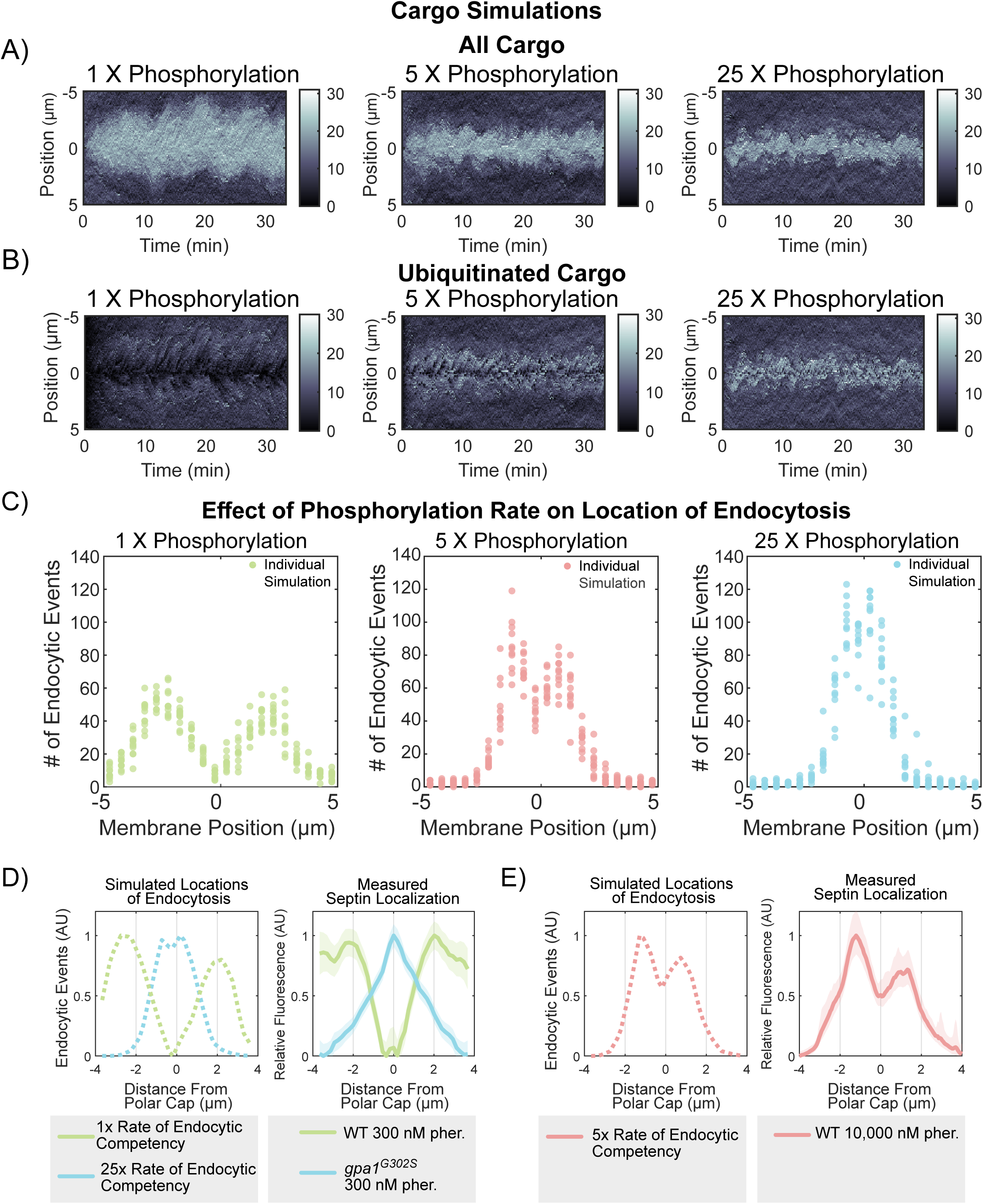
Computational modelling predicts that endocytosis dynamics are capable of generating septin distribution patterns. To change endocytic competency rate, we modified the phosphorylation probability 1-, 5-, and 25-fold. A) Kymographs from individual simulations showing all cargo on the membrane over time. B) Kymographs from individual simulations showing ubiquitinated cargo on the membrane over time. C) Histograms of the location of endocytic events for 11 independent simulations per phosphorylation probability (1-, 5-, and 25-fold). 1X phosphorylation rate yields a distribution of endocytic events that mirrors WT septin localization. While increased phosphorylation probabilities (5-fold and 25-fold higher) result in endocytic events occurring closer to the site of polarity. D) Comparisons of the distribution of simulated endocytic events with experimentally determined septin distributions. All plots have been normalized to have a maximum value of 1. Comparison of the average location of endocytic events (from C) with low (1x phosphorylation) and high (25x phosphorylation) rates of endocytosis competency from Figure 6 with the WT and *gpa1^G302S^* septin distributions in 300 nM pheromone from Figure 1. E) The model suggests that a higher rate of endocytosis competency would push endocytic events closer to the center of the polar cap (on the left). Since we hypothesize that the rate of endocytosis competency is influenced by Gpa1 activity levels, we added a high level of pheromone (10 µM) and measured septin distribution (right). Shaded area represents 95% confidence intervals, n = 293 cells across two independent experiments.

These changes also influenced the spatial location of endocytic events, which moved closer to the center of the polar cap at higher phosphorylation rates (Figure 7C). The accumulation of ubiquitinated cargo near the center promoted the formation of new endocytic sites in this region, reinforcing the centralization of endocytosis. The increased phosphorylation rates effectively reduced the time cargo could diffuse along the membrane, limiting its ability to travel farther from the polarity site before being internalized.

When we alter the rate that cargo becomes competent for endocytosis in our simulations, we find that the location of endocytic events qualitatively recreates the septin distributions we have measured in WT and *gpa1^G302S^*strains at 300 nM pheromone. When we compare our simulation of endocytosis locations to our experimentally measured septin distributions, we see that simulations of 1X phosphorylation rate leads to endocytic events with a pattern similar to septin distribution in our WT cells treated with 300 nM pheromone. The 25X phosphorylation rate simulations generate endocytic events with a distribution similar to experimental septins in the hyperactive *gpa1^G302S^* mutant strain (Figure 7D).

Our simulations suggest that a 5x rate of phosphorylation would lead to endocytosis events closer to the polar cap than we had measured at 300 nM pheromone (Figure 7C). If we are correct that the location of endocytosis is dictating the location of septin organization and our model is accurately recapitulating the system, then we would expect that the endocytic distribution of the 5X phosphorylation rate would match the septin distribution in a WT cell in the presence of a high dose of pheromone. In this case, the pathway activity would be high, but the Gα would still be able to turn off. We treated WT cells with 10 µM pheromone (a saturating dose) for 2 hours and measured Cdc3-mCherry distribution relative to the polar cap (Figure 7E). The measured septin distribution matches the simulation remarkably well, even down to the same asymmetry in both simulated endocytic events and septin distribution, which we attribute to chance, as we have no reason to believe there is an underlying asymmetry in the biology and our model should not have any intrinsic asymmetry. These data do not necessarily mean that *gpa1^G302S^* increases the phosphorylation of a cargo per se, but it does suggest that increasing the rate that a cargo becomes competent for endocytosis and coupling that endocytic event to the assembly of macromolecular complexes of septin would be capable of creating the septin distributions we see in WT and *gpa1^G302S^* mutants.

### Clathrin-mediated endocytosis is required for Gpa1 control of septin organization

With our model indicating that endocytosis can generate the spatial patterns we see with septins and our data implicating Ent1 and Ent2 in septin regulation, we hypothesized that disruption of clathrin-mediated endocytosis would also rescue septin organization in the *gpa1^G302S^* mutant strain. To disrupt endocytosis, we deleted End3, an endocytic protein that works in a separate complex from Ent1 and Ent2 and is required for clathrin-mediated endocytosis (Figure 8A) (Benedetti et al., 1994; Sun et al., 2015). Deletion of *END3* from cells with WT GPA1 lead to broader projections, but minimal impact on the relative localization of septins (Figure 8B). Deletion of *END3* diminishes the already low frequency of class 1 septin usage (Figure 8D). Deletion of *END3* from *gpa1^G302S^* cells leads to a rescue of septin localization to the polar cap based on the average distribution (Figure 8C). When analyzing septin class usage, this rescue happens through an increase in usage of the asymmetric classes 4, 5, and 6 rather than an increase in class 2. Nonetheless, when we examine the similarity of septin usage to WT *GPA1* and *gpa1^G302S^*, End3 deletion leads to an overall septin usage that is closer to WT than *gpa1^G302S^*(Figure 8E).

**Figure 8.**
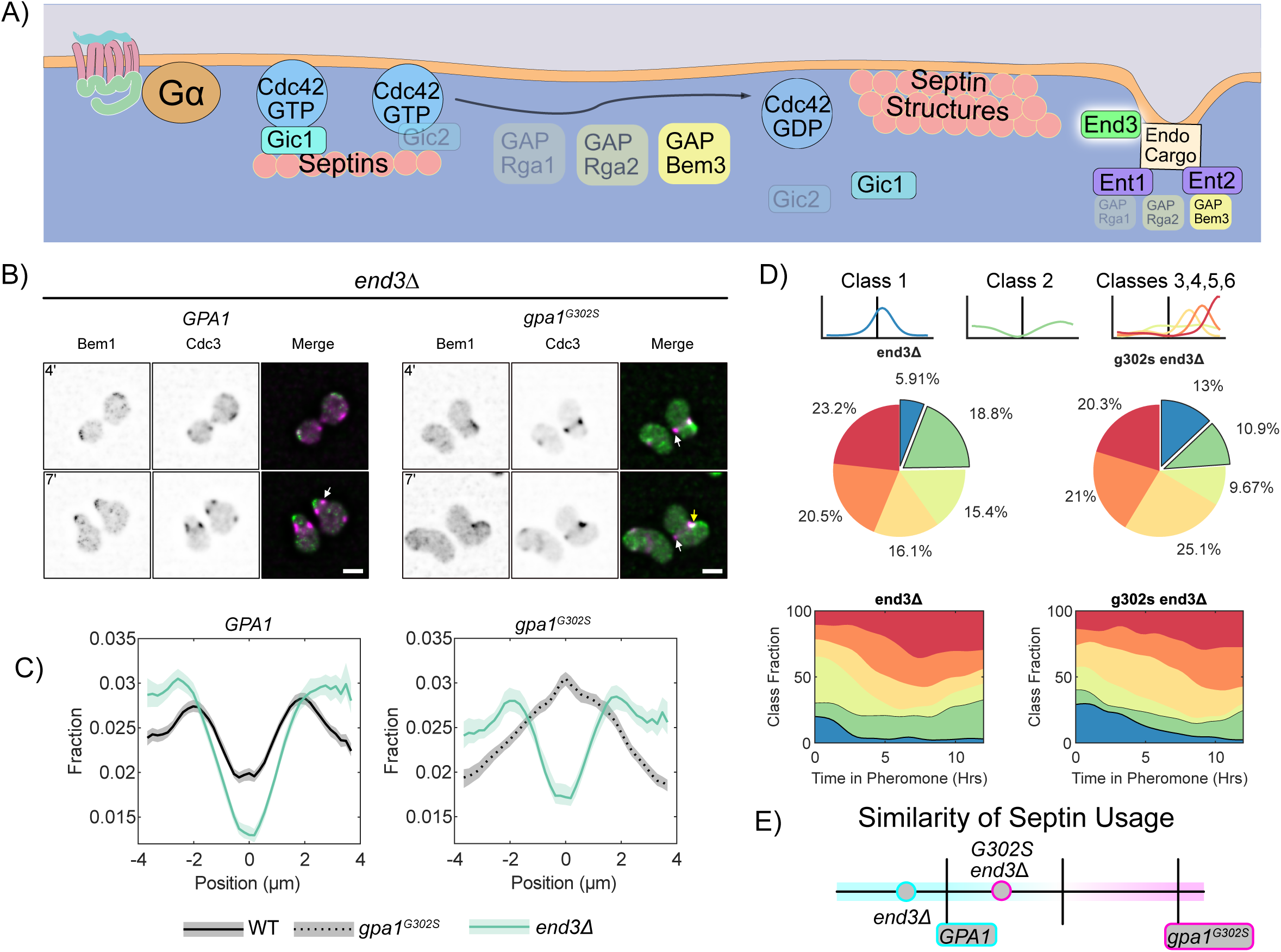
The Clathrin Mediated Endocytosis protein End3 is required for Gpa1 control of septin organization. A) Schematic showing the role of End3 as an endocytic protein that interacts with the epsins Ent1 and Ent2. B) Representative images of *end3*Δ *BEM1-GFP CDC3-mCherry* cells with WT *GPA1 or gpa1^G302S^* at 4 and 7 hours of treatment with 300nM pheromone. Scale bar represents 5µm. C) Average septin profiles from 12-hour microfluidic experiments, normalized to the center of the polar cap, for each of the indicated strains in *GPA1* or *gpa1^G302S^*backgrounds. WT *GPA1* and *gpa1^G302S^*without further gene deletions are from figure 1 and reproduced for ease of comparison. Shaded area represents 95% confidence intervals. Lines are representative of at least 2 combined experiments: *end3*Δ n= 50 cells, *gpa1^g302s^ end3*Δ n = 50 cells. D) Septin class usage in end3Δ cells, with and without the *gpa1^G302S^*mutant, overall (pie charts) and over time (stacked graph). E) Similarity of the septin usage for each deletion strain compared to WT *GPA1* or *gpa1^G302S^* alone based on septin class frequency distance calculation.

### Endocytosis of Gα and the pheromone receptor directs pheromone-induced septin distribution

Having established that endocytosis plays a central role in Gpa1 control of septin organization, we set out to find the relevant cargo. The most well-characterized endocytic cargo during the pheromone response is the receptor, Ste2 (Figure 9A), which is both phosphorylated and ubiquitinated at its C-terminus to control its endocytosis (Dunn and Hicke, 2001; Hicke et al., 1998). We constructed cells lacking the receptor C-terminus from codon 326 (*ste2*^Δ*CT*^), a critical region for control of receptor endocytosis (Konopka et al., 1988). In WT Gpa1 cells, deletion of the C-terminus of the receptor had no obvious effect, but when we look at class usage, we see a marked increase in class 2, where septins are symmetrically distributed around the polar cap. We then investigated the effect of the *ste2*^Δ*CT*^ on septin localization in cells expressing *gpa1^G302S^*. We found that *gpa1^G302S^*mutants lacking the C-terminus of Ste2 showed a small change in septin organization (Figure 9B, D, E). Morphologically, the *gpa1^G302S^ ste2*^Δ*CT*^ double mutants are very round, with poor mating projection formation, consistent with septin defects. Comparing the similarity of septin usage (Figure 9G), *ste2*^Δ*CT*^ is much closer to *gpa1^G302S^* than WT. Thus, in the presence of WT Gpa1, the C-terminus of the receptor can drive an increase in asymmetric septin usage, indicating that it may play a role in normal septin organization, but is not dominant to the gpa1^G302S^ mutant. Thus, there must be another cargo driving septin distribution.

**Figure 9.**
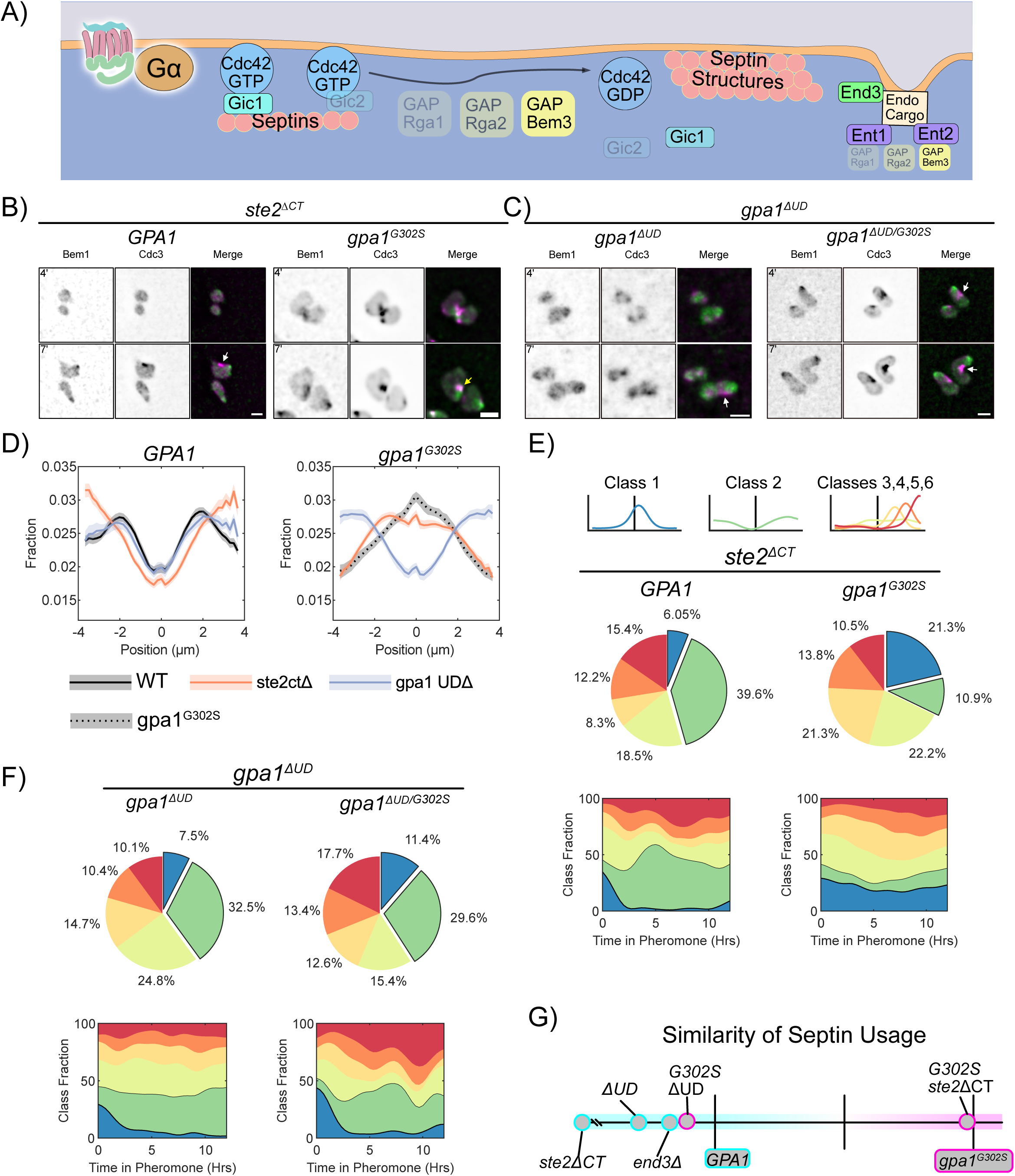
Gα control of septin deposition is driven by its ubiquitination domain. A) Schematic showing known endocytic cargoes, Gpa1 and Ste2. B) Representative images of *ste2*^Δ*CT*^ *BEM1-GFP CDC3-mCherry* cells with WT *GPA1 or gpa1^G302S^*at 4 and 7 hours of treatment with 300nM pheromone. Scale bar represents 5µm. C) Representative images of *BEM1-GFP CDC3-mCherry* cells with *gpa1*^Δ*UD*^ *or gpa1*^Δ*UD/G302S*^ at 4 and 7 hours of treatment with 300nM pheromone. Scale bar represents 5µm. D) Average septin profiles from 12-hour microfluidic experiments, normalized to the center of the polar cap, for each of the indicated strains with or without the *gpa1^G302S^*mutation. WT *GPA1* and *gpa1^G302S^* without further gene deletions are from figure 1 and reproduced for ease of comparison. Shaded area represents 95% confidence intervals. Lines are representative of at least 2 combined experiments: *ste2*^Δ*CT*^ n= 58 cells, *gpa1^g302s^ ste2*^Δ*CT*^ n = 140 cells, *gpa1*^Δ*UD*^ n = 50 cells and *gpa1*^Δ*UD/G302S*^ n = 118 cells). E) Septin class usage in *ste2*^Δ*CT*^ cells, with and without the *gpa1^G302S^* mutant, overall (pie charts) and over time (stacked graph). F) Septin class usage in *gpa1*^Δ*UD*^ *or gpa1*^Δ*UD/G302S*^ mutants, overall (pie charts) and over time (stacked graph). G) Similarity of the septin usage for each deletion strain compared to WT *GPA1* or *gpa1^G302S^* alone based on septin class frequency distance calculation.

Gpa1 contains a ubiquitination domain (UD) that promotes endocytosis and subsequent trafficking through the endomembrane system (Dixit et al., 2014a; Wang et al., 2005a). We hypothesized that endocytosis of Gpa1 (via the UD) directs septin deposition. We deleted the UD from both *GPA1* and *gpa1^G302S^*. Cells expressing gpa1 ^Δ^*^UD^* without the G302S substitution display an average septin organization similar to WT (Figure 9D). However, much like the *ste2*^Δ*CT*^ strain, they showed a marked increase in the usage of class 2 septins (Figure 9F).

When we examined *gpa1^G302S-^*^Δ*UD*^ cells exposed to 300nM pheromone – as all other strains were exposed to – we found that cells reverted to mitotic behavior after only 8 hours of pheromone exposure (Movie S1). While WT cells will sometimes revert to mitotic behavior in the microfluidic device on this time scale, the supersensitive *gpa1^G302S^*never do. While the *gpa1^G302S^* mutant increases pathway sensitivity, the *gpa1*^Δ*UD*^ mutant decreases pathway sensitivity (DiBello et al., 1998; Wang et al., 2005b). The deletion of the UD, therefore, counteracts the supersensitivity due to reduced RGS interaction from the G302S mutation, at least for cell cycle arrest. For the purposes of our septin experiments, we found that increasing the concentration of pheromone to 1µM lead to the *gpa1^G302S-^*^Δ*UD*^ cells continuing to develop mating projections for the full 12-hour time course (Movie S1), although both doses yield the same septin distributions (Figure S2).

Remarkably, deletion of the UD from gpa1^G302S^ was sufficient to rescue septin organization. Cells expressing *gpa1^G302S-^*^Δ*UD*^ displayed septin structures well separated from the polar cap (Figure 9C). This rescue was evident in the quantitation of both the average septin profile and in the analysis of septin class usage. Specifically, there was a marked decrease in the proportion of class 1 septins, accompanied by a robust increase in the use of class 2 septin shapes, and a notable rise in class 6 (Figure 9F). Furthermore, similarity analysis of septin usage (Figure 9G) demonstrated that UD deletion restored septin usage to a pattern similar to WT cells, whereas deletion of the receptor c-terminus had minimal impact. These findings indicate that Gpa1 control of septins depends upon its UD.

We have previously shown that the septin defects generated by *gpa1^G302S^*are not due to pathway sensitivity, as *sst2^Q304N^* shows the same pheromone sensitivity as *gpa1^G302S^*, but does not drive septin colocalization with the polar cap (Dixit et al., 2014b; Kelley et al., 2015). Nonetheless, given that *gpa1^G302S-^*^Δ*UD*^ affects pathway sensitivity, we tested the sensitivity of the *gpa1^G302S^* background strains using a halo assay (Hoffman et al., 2002). We find that all other *gpa1^G302S^*background strains retain supersensitivity (Supplemental Figure S3), confirming that the rescue of septin organization is not simply due to altered pathway sensitivity.

These results demonstrate that Gpa1 controls septin organization through its UD. Interestingly, Ste2 influences septin organization via its C-terminus through a parallel pathway.

### Analysis of septins usage across strains

The extensive septin data generated in this project allowed us to develop a comprehensive understanding of the impacts of these gene deletions on septin organization. We determined the distance between the septin class frequency data for each strain and performed hierarchical clustering (Figure 10A). We performed bootstrapping on the hierarchical clustering, but only 4 groupings reached 95% confidence. These 4 groups, highlighted in the blue region, cluster closely with WT septin usage.

**Figure 10.**
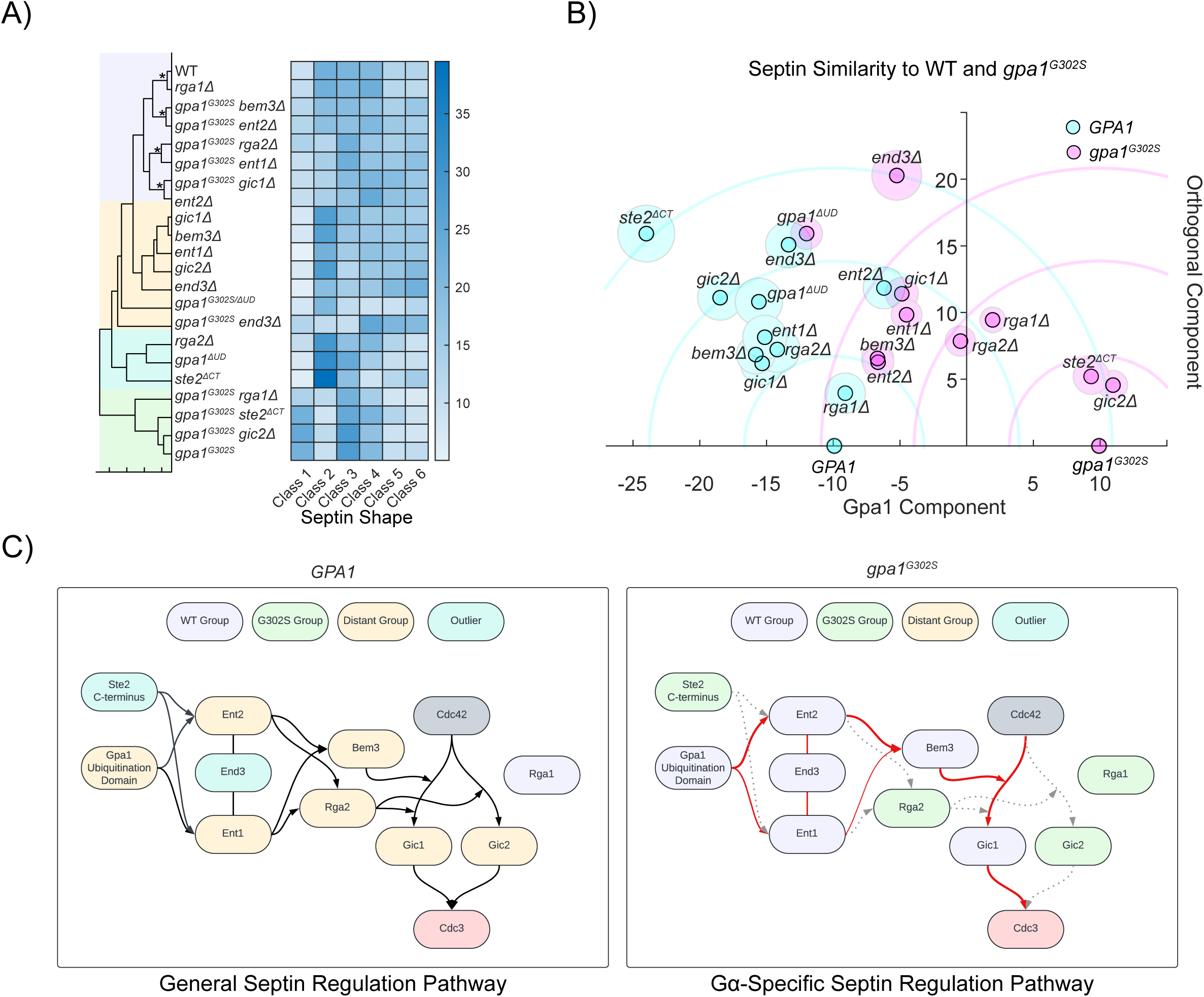
Analysis of septin usage throughout this study reveals the pathway used for Gpa1 control of septin organization. A) Hierarchical clustering of each strain in this study based on their septin class usage (72,222 different septin profiles across the entire study). Shaded boxes represent groups referred to in part C. Strains within the same nodes that achieved ≥95% confidence by bootstrapping are marked with *. B) This graph is a projection of the distance between each strain (based on septin usage) onto a coordinate plane defined by an X-axis containing the WT and gpa1^G302S^ strains. Each strain is then mapped onto the positive Y-axis based on its distance from WT and gpa1^G302S^. This can be thought of as the intersection of two circles, one with a radius defined by the distance from WT, and the other defined by the distance from mutant Gpa1. Shaded circles represent 95% confidence intervals derived through bootstrapping. Each strain’s X-value from this coordinate system is reported throughout this study for the “Similarity of Septin Usage” (Figures 3G, 4F, 5F, 7E, 8G). C) Map of the signaling pathway that controls septin organization in the presence of WT Gpa1, and in the presence of Gpa1^G302S^. In this epistasis analysis, genes are considered to be part of the general septin regulation pathway if septin usage changes when the gene is deleted from an otherwise WT cell, and to be part of the Gα-specific septin regulation pathway if septin regulation is closer to WT than *gpa1^G302S^* when the gene is deleted in a *gpa1^G302S^* background.

We have the advantage of having both the *GPA1* and *gpa1^G302S^* backgrounds for every deletion strain, enabling a comparative distance analysis between these genotypes, which differ only in the activity level of Gpa1. By defining a coordinate plane where the X-axis represents the distance between *GPA1* and *gpa1^G302S^*, each additional strain can be placed on the coordinate plane based on their respective distances. Conceptually, this analysis is the intersection of two circles, though we consider only the positive intersection values(see Figure 10B). We performed bootstrapping to establish 95% confidence intervals for the position of each strain, yielding a clear representation of the organization of these strains.

The X-axis, or “Gpa1 axis,” quantifies each strain’s similarity to GPA1 or *gpa1^G302S^*, providing a metric for the Similarity of Septin Usage that we have reported above (Figures 3G, 4G, 5G, 8E, 9G). The Y-axis is an orthogonal component that reflects other unknown aspects of septin regulation. It is noteworthy that the strains that have the highest values on this orthogonal axis are *the Gpa1 and Ste2 endocytosis-incompetent strains end3*Δ*, gpa1^G302S^ end3*Δ*, gpa1^G302S-^*^Δ*UD*^ and *ste2*^Δ*CT*^, suggesting some link between endocytosis and this dimension. Notably, *gpa1^G302S^ bem3*Δ and *gpa1^G302S^ ent2*Δ have almost identical locations on this graph, indicating that the Bem3 and Ent2 may contribute to septin organization during the pheromone response through their interaction with each other.

To further explore these relationships, we performed an epistasis analysis using the clustering data. We reasoned that gene deletions that disrupt septin organization in a *GPA1* background must contribute to the general regulation of septins during the pheromone response. In contrast, genes that restore WT-like septin organization in the *gpa1^G302S^* background must specifically contribute to Gpa1 control of septins. We integrated this analysis with known protein interactions to generate the diagrams in Figure 9C. We observed that RGA1 plays virtually no role in septin organization during the pheromone response, regardless of Gpa1 involvement, while all other genes examined play a role in the general regulation of septins. Notably, Gpa1 specifically relies upon a pathway that goes from the endocytic proteins End3, Ent1, and Ent2, through the Cdc42 regulator Bem3 and Cdc42 effector Gic1 to regulate septins.

## Discussion

Here, we describe a mechanism by which Gpa1 regulates septins during the yeast pheromone response. By systematically investigating the roles of known septin regulators, we identified Gic1 as a critical effector, while its paralog Gic2 was dispensable. We found that the Cdc42 GAP Bem3 preferentially mediates this process, rather than Rga1 or Rga2. Given the established interactions between the Cdc42 GAPs and the endocytic adaptors Ent1 and Ent2, we tested whether these epsins contribute to Gpa1-regulated septin organization. We found that Ent1 and Ent2 function redundantly to promote septin organization in response to pheromone. This led us to hypothesize that endocytosis was providing spatial cues for septin organization. We developed an agent-based mathematical model to test if the location of endocytic events could explain septin distribution. The model predicts endocytic event locations that recapitulate the septin distributions we see in wild-type and mutant Gpa1 contexts if we assume that Gpa1^G302S^ increases the rate at which cargo is marked for clathrin-mediated endocytosis. Experimental validation confirmed that clathrin-mediated endocytosis, disrupted by END3, impacts septin organization. While we hypothesized that endocytosis of the pheromone receptor Ste2 might mediate Gpa1 control of septins, disrupting Ste2 endocytosis had only a mild effect in the presence of gpa1^G302S^. The Ste2 C-terminus did, however, regulate septin organization in the presence of WT Gpa1, indicating a parallel pathway that is recessive to the effects of gpa1^G302S^. Gpa1 internalization is less well characterized than Ste2 but is known to be dependent upon its Ubiquitin Domain (UD) (Dixit et al., 2014a). Remarkably, blocking Gpa1 endocytosis by removal of the UD led to the complete recovery of septins to the periphery of the polar cap in the presence of the hyperactive *gpa1^G302S^*mutation. Thus, Gpa1 controls septins through a mechanism involving its UD, clathrin-mediated endocytosis machinery (including End3, Ent1, and Ent2), the Ent2-interacting Cdc42 GAP Bem3, and Gic1.

### The Endocytic Machinery as a Regulator of Septin Organization

Septin organization occurs during mitosis independently of Gα activity. Similarly, we demonstrated that Gpa1 activity is dispensable for septin organization during the pheromone response, as signaling initiated by the MAPK scaffold Ste5 is sufficient to drive septin structure formation at the base of the mating projection. Thus, Gpa1 introduces an additional layer of septin regulation on top of the normal process.

Our data are consistent with the following model (Figure 11): In the presence of pheromone, phosphorylation and subsequent ubiquitination of Gpa1 and Ste2 facilitate their recruitment of epsins and the Cdc42 GAP Bem3 to sites of endocytosis. The recruitment of Bem3 promotes Cdc42 hydrolysis of GTP and subsequent organization of higher-order septin structures in a Gic1-dependent manner. When Gpa1 is hyperactive, these events happen closer to the polar cap. We speculate that Gpa1^G302S^ becomes competent for endocytosis earlier, speeding the recruitment of epsins and Cdc42 GAPs. Our ability to monitor Gpa1 trafficking is limited, as neither N- nor C-terminal fusions of GFP with Gpa1 are functional. Future studies will be required to test this possible mechanism linking Gpa1 activation state to its trafficking.

**Figure 11.**
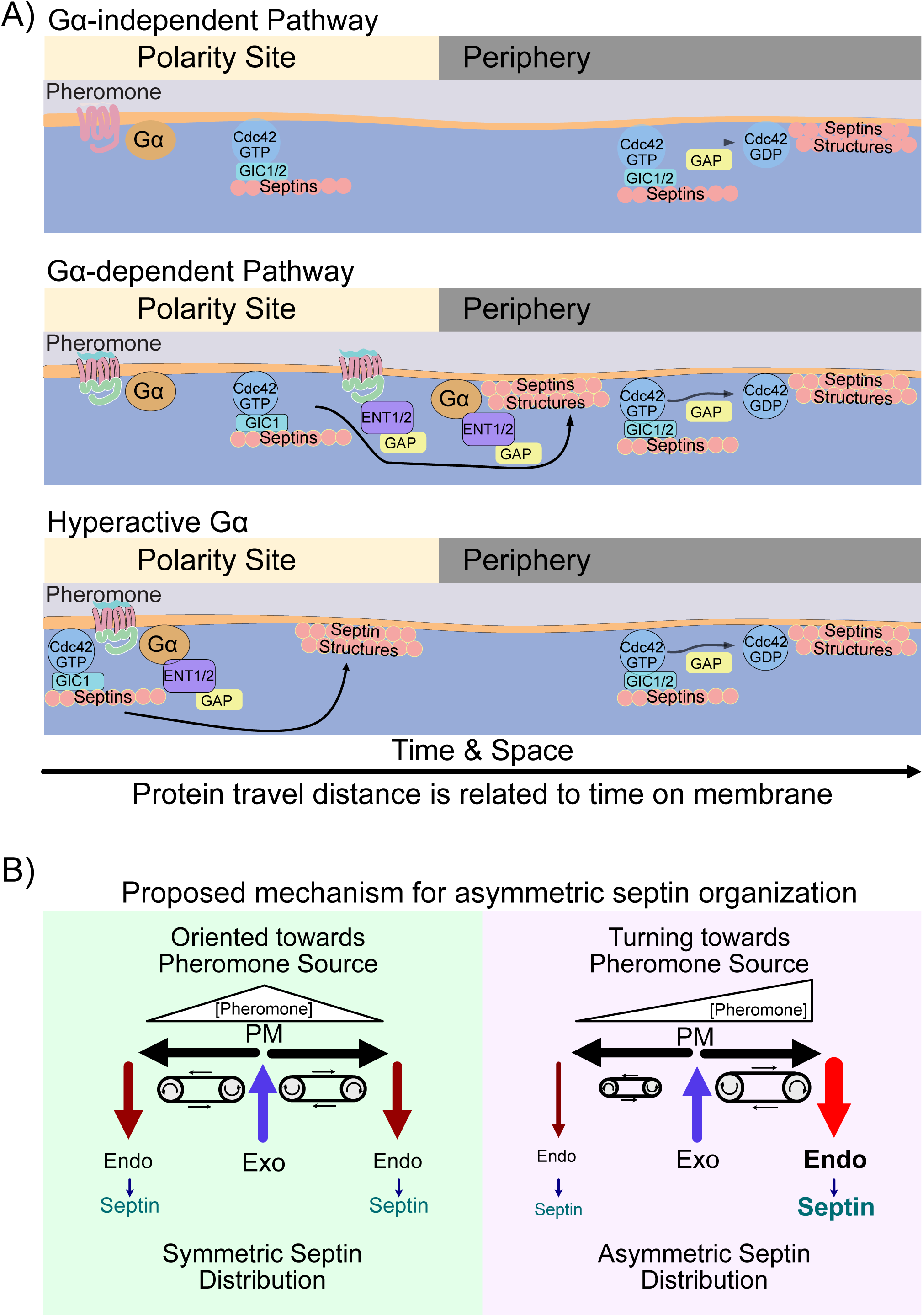
Diagram of proposed mechanism. A) Proposed molecular mechanism for three scenarios: 1) The Gpa1 independent pathway, which would use Gics and Cdc42-GAPs to promote septin organization, similar to mitotic processes. 2) The standard Gpa1-regulated pathway, where we propose Gpa1 recruits epsins and Cdc42-GAPs to sites of high Gpa1 activity, leading to septin organization. The receptor C-terminus also appears to regulate septins under these conditions. and 3) The Hyperactive Gpa1 scenario, where Gpa1 activity is high close to the center of the polar cap, leading to faster recruitment of endocytic machinery, and therefore septin organization closer to the center of the polar cap than normal. Consistent with our mathematical modeling, proteins that stay on the membrane longer also travel further from the polar cap, thus faster recruitment of endocytic adaptors leads to less distance traveled. B) Proposed mechanism for the asymmetric distribution of septins during gradient tracking. Note that delivery of vesicles to the center of the polarity site and removal distal to the polarity site creates a “treadmill” effect where plasma membrane moves away from the center and towards the sites of endocytosis. On the left, when the polar cap (where exocytosis is occurring) is oriented towards the source of pheromone, the amount of signal to either side of the polar cap is equal, Gpa1 activation levels are similar, leading to similar amounts of endocytosis, and symmetric septin organization. On the right, the polarity site (where exocytosis is occurring) is not oriented towards the source of pheromone, instead high pheromone to the right of the polarity site is driving increased endocytosis relative to the opposite side of the polarity site, leading to asymmetric distribution of septins.

Previous models of septin organization incorporated exocytosis as a driving force for clearance of septins from the bud site, and Cdc42 GAP function as the “timer” dictating how far septins were pushed before assembly into higher order structures (Okada et al., 2013). Our findings further this framework by suggesting that epsin-mediated endocytosis may control *where* septins transition into higher-order structures. Thus, vesicle trafficking emerges as a dual driver in this process, moving septins away from the polar cap on a “conveyor belt” of the plasma membrane (moving away from exocytic events and towards endocytic events) (Figure 9B). Coupled to this, the endocytic machinery provides the spatial cue that septins should transition from “passengers” on this conveyor belt to establishing higher-order structures.

In the absence of the hyperactivating G302S mutation in GPA1, disruption of endocytosis of both Ste2 and Gpa1 lead to a large increase in the frequency of symmetric septin organization (class 2), indicating that these domains promote asymmetric septin organizations. The observation that both proteins influence septins in this way raises the possibility that endocytic cargoes, in general, may play a broader role in septins organization. The fact that Ste2 C-terminal deletion was recessive to the G302S mutation in GPA1 indicates that they work upon parallel pathways to regulate septins, which further supports the idea that each can influence septin organization through their endocytosis. Both receptor and Gα undergo endocytosis in response to pheromone, but Ste2 endocytosis is likely not linked to Gpa1 activation.

Our model for Gα control of septins during the pheromone response predicts that the distance between the polar cap and septin structures decreases with higher pheromone doses and increases at lower doses. Consistent with this, we found that septins accumulated closer to the polarity site at 10 uM pheromone than at 300nM pheromone (Figure 7B). Similarly, in our previous work (Kelley et al., 2015), we found that cells responding to an intermediate of pheromone dose (∼75 nM, center of a 0-150 nM gradient) established septin structures farther from the polar cap center than cells exposed to a high dose of pheromone (300 nM). These findings align with our model, which couples pheromone dose to septin spatial regulation, which is necessary for a mechanistic understanding of gradient tracking.

### Implications for gradient tracking

We previously found that cells without functional septins grow in a straight line, unable to correct their orientation by turning (Kelley et al., 2015). WT cells that turn display septin enrichment on the inside of the turn. Gpa1G302S mutant cells likewise turn towards their mislocalized septins. Thus, the asymmetric distribution of septins on the inside of the turn (up-gradient) appears to be required for error correction during gradient tracking (Kelley et al., 2015). Our findings that Gpa1 activity level influences septin organization through the endocytosis machinery provides a potential mechanism for asymmetric septin organization (Figure 9B). Specifically, our data supports a model where higher active Gpa1 on the up-gradient side of the cell results in increased Gpa1 endocytosis. The associated endocytic machinery promotes septin organization on the up-gradient side of the cell. This accumulation of septins could then facilitate the turning of the cell. Future studies will be needed to address how Gpa1 activity influences its internalization through the UD.

### Epsins and Cdc42-GAPs

Our findings raise the question of whether endocytic events also control the organization of the septin ring during mitosis. Fortunately, the Aguilar lab has provided evidence supporting this idea. They demonstrated that overexpression of the Epsin N-terminal Homology Domain from Ent2 (ENTH2) leads to defects in the organization of septins during mitosis (Mukherjee et al., 2009). Furthermore, they found that ENTH2 overexpression affects Bem3 localization, suggesting that Epsin recruitment of Bem3 may be a broadly used mechanism for controlling septin organization.

Separately, the Bi and Goryachev labs showed that during mitosis, exocytosis drives septin formation into a ring structure, with GAP activity proposed to limit ring expansion. This mechanism results in septin organization at the edge of the polar cap (Okada et al., 2013). The initial cue for septin structure formation during mitosis may arise from mechanisms distinct from exocytosis, such as endocytic cargoes recruiting Epsin/Bem3 complexes. Subsequent stabilization of these structures might then involve Rga1 and Rga2, which are known to bind to mitotic septins and could reinforce their formation. By contrast, Rga1 and Rga2 show little localization during the pheromone response (Kelley et al., 2015). This observation and our new findings suggest that Bem3 plays the primary role in controlling septins during mating. These differences underscore distinct regulatory mechanisms for septin organization during mitosis versus the pheromone response, with Epsin/Bem3 complexes playing a potentially pivotal role in both contexts.

### Conservation of Epsins and G-protein endocytosis from yeast to humans

We have demonstrated that Gpa1 influences septin organization through its unique yeast-specific UD, which is required for Gpa1 endocytosis (Figure 9) (Dixit et al., 2014a; Wang et al., 2005b). The molecular mechanisms underlying the endocytosis of Ste2 and Gpa1 are unique to yeast, including the presence of the UD, which is specific to yeast Gα proteins (Wang et al., 2005b). Unlike in yeast, mammalian Gα endocytosis occurs indirectly via receptor C-terminal interactions, rather than through direct internalization of the Gα subunit (Shenoy and Lefkowitz, 2003; Smith et al., 2021). Despite these species-specific differences in GPCR and Gα internalization, the role of epsins is well conserved from yeast to humans.

Epsins are central to clathrin-mediated endocytosis, serving as cargo adaptors and facilitating vesicle formation. Their functional conservation is evident in mammalian systems, where they interact with AP2 (Chen et al., 1998), bind Cdc42-GAP (Coon et al., 2010), and contain a ubiquitin-interacting motif essential for cargo recognition (Wendland, 2002). Epsins also promote the endocytosis of human proteins, such as the epidermal growth factor (EGF) receptor (Kazazic et al., 2009). Thus, there may be a conserved mechanism to couple septin organization to endocytosis across species.

## Abbreviations

GPCR: G-protein Coupled Receptor
UD: Ubiquitination Domain
CT: C-terminus
GAP: GTPase Activating Protein
RGS: Regulator of G-protein Signaling

## Acknowledgements

The authors wish to thank Holly Goodson, Melissa Maginnis, and Amy Gladfelter for their helpful discussions. The authors would also like to thank Niklas Hase, the Shmoo Crew and other former members of the Kelley Lab for their help and input. This work was supported by the National Institutes of Health Grants R15-GM128026 to JBK and AK, and R15-GM140409, R15-GM155864, and P20-GM144265 to JBK.

## Author Contributions

CPJ wrote the manuscript -original draft, review, and edits, visualization; designed, conducted, analyzed, and oversaw the experiments. SS wrote the manuscript -review and edits, collected and analyzed data. AH wrote the manuscript -review and edits, collected, and analyzed data. KJF wrote the manuscript -original draft, review, and edits, designed and conducted experiments (mathematical modeling), and created software. LG wrote the manuscript – review and edits, collected and analyzed data. SGL wrote the manuscript -review and edits, collected, and analyzed data. NL wrote the manuscript -review and edits, collected and analyzed data. RG wrote the manuscript - review and edits, analyzed data. TS wrote the manuscript -review and edits, collected and analyzed data. MS wrote the manuscript -review and edits, collected and analyzed data. AK wrote the manuscript-review and edits, and contributed supervision. JBK wrote the manuscript -original draft, review and edits, visualization, designed experiments, acquired funding, contributed supervision, conceptualized and oversaw the project.

