## Supplemental FIgures for "Septin organization is regulated by the Gpa1 Ubiquitination Domain and Endocytic Machinery during the yeast pheromone response"

### Supplemental Figure S1

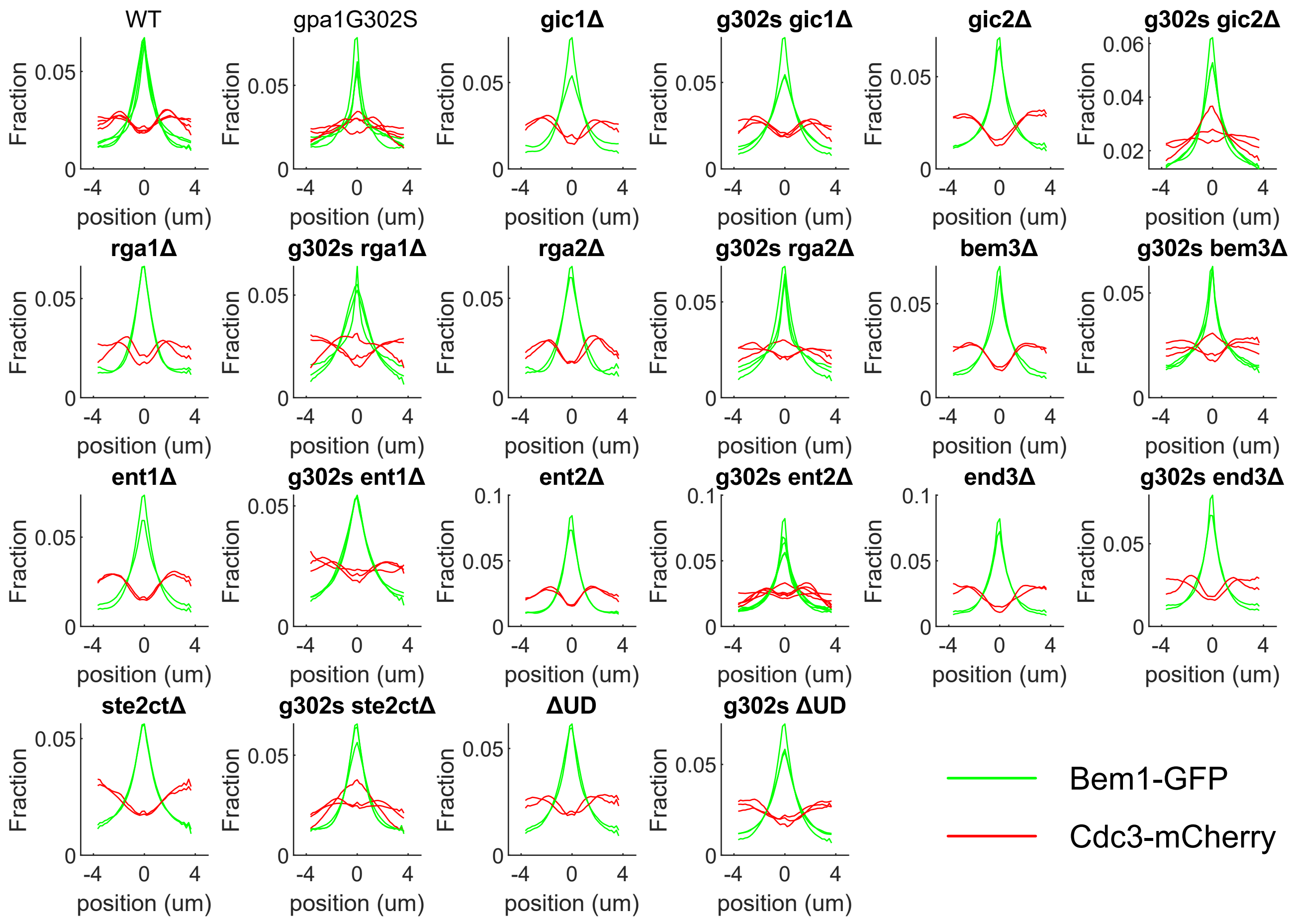


**Supplemental Figure S1. Experiment-to-experiment Variability.** The average Bem1 (green) and Cdc3 (red) profiles for each independent experiment for the indicated strain are shown. Strains with *gpa1*$^{G302S}$ backgrounds show higher experiment-to-experiment variability.

### Supplemental Figure S2

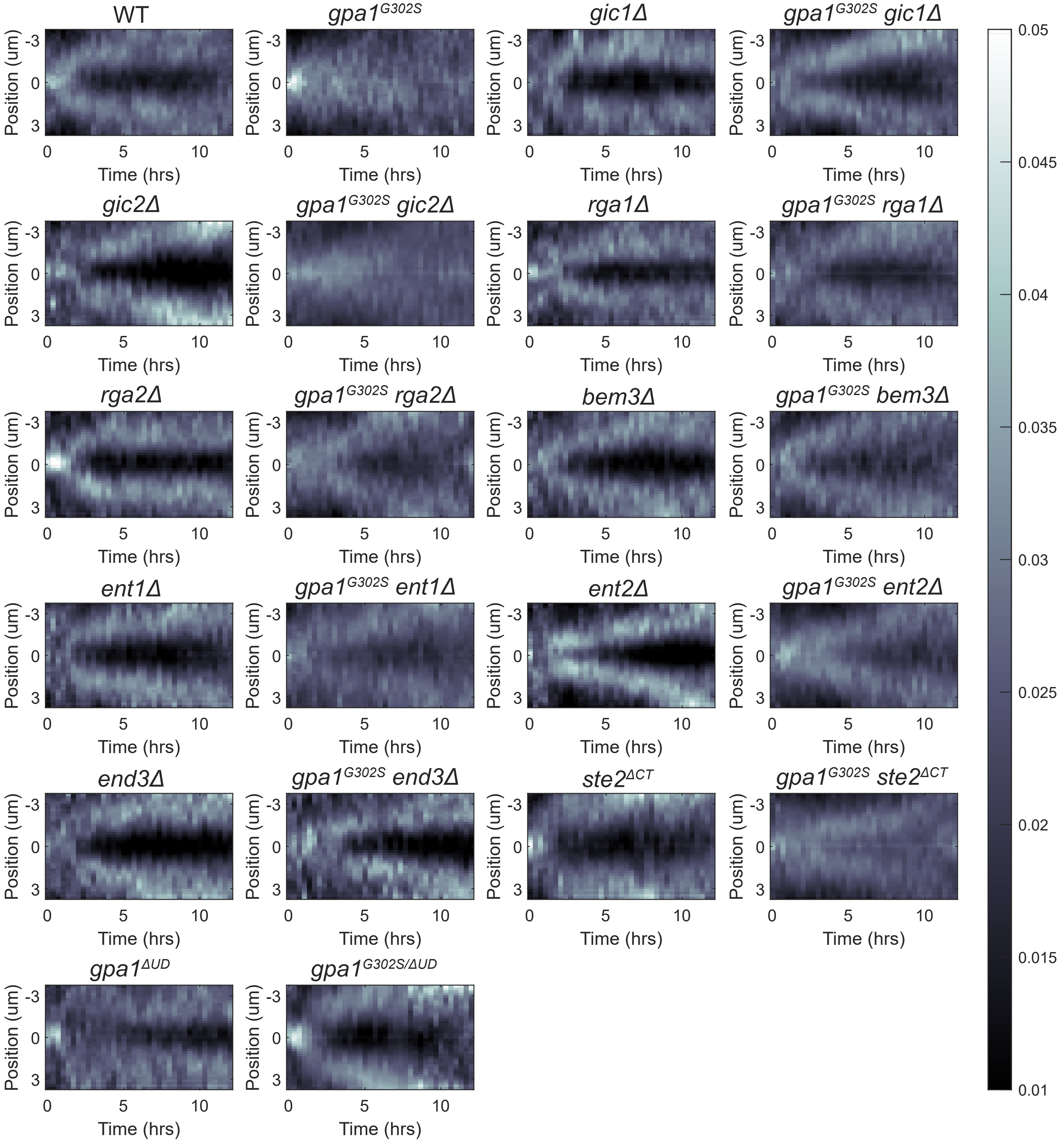


**Supplemental Figure S2. Average Kymographs of Cdc3-mCherry.** The average kymograph of Cdc3-mCherry spatially normalized to peak Bem1-GFP signal for every cell from the indicated strain.

### Supplemental Figure S3

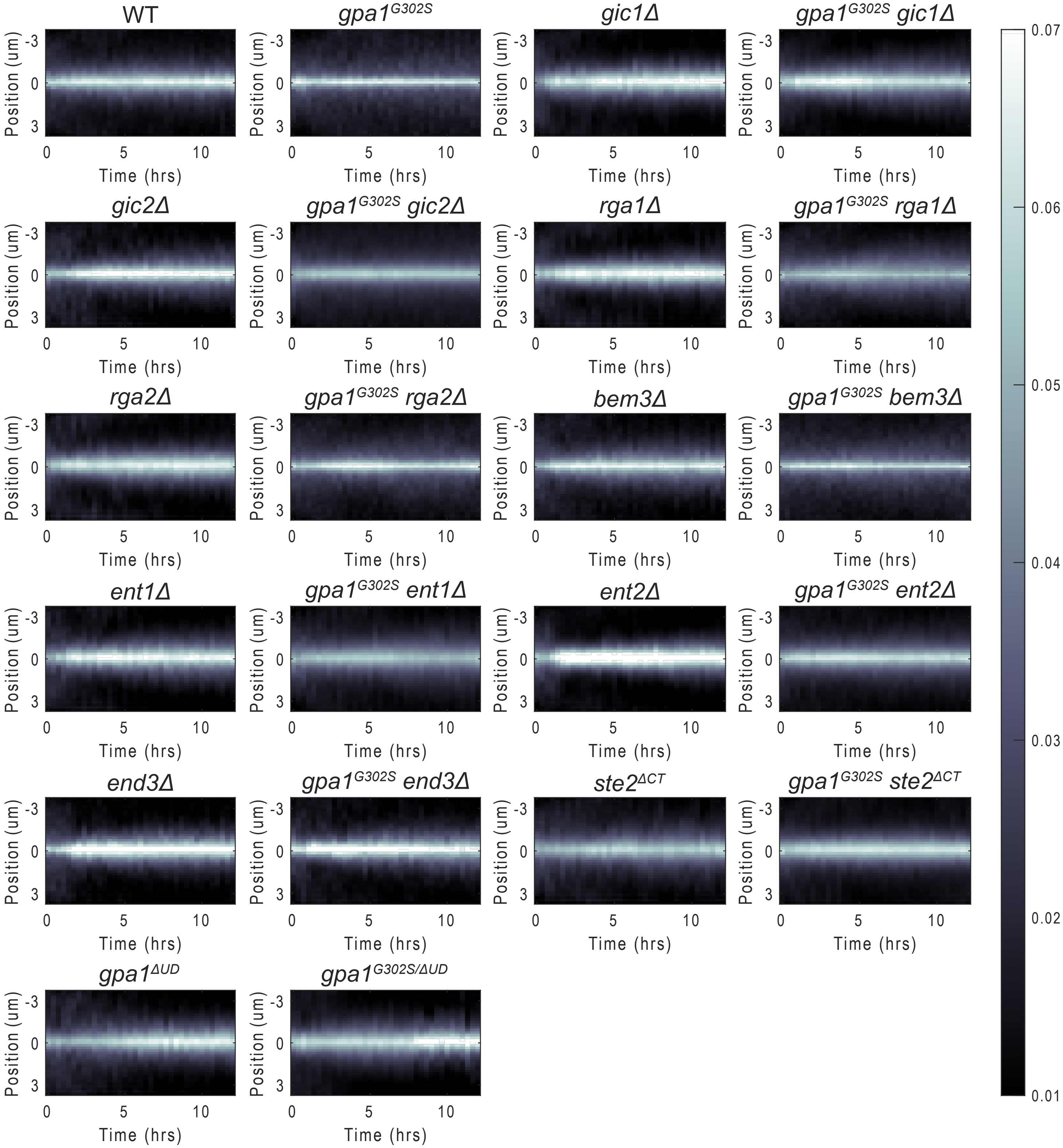


**Supplemental Figure S3. Average Kymographs of Bem1-GFP.** The average kymograph of Bem1-GFP spatially normalized to peak Bem1-GFP signal for every cell from the indicated strain.

### Supplemental Figure S4

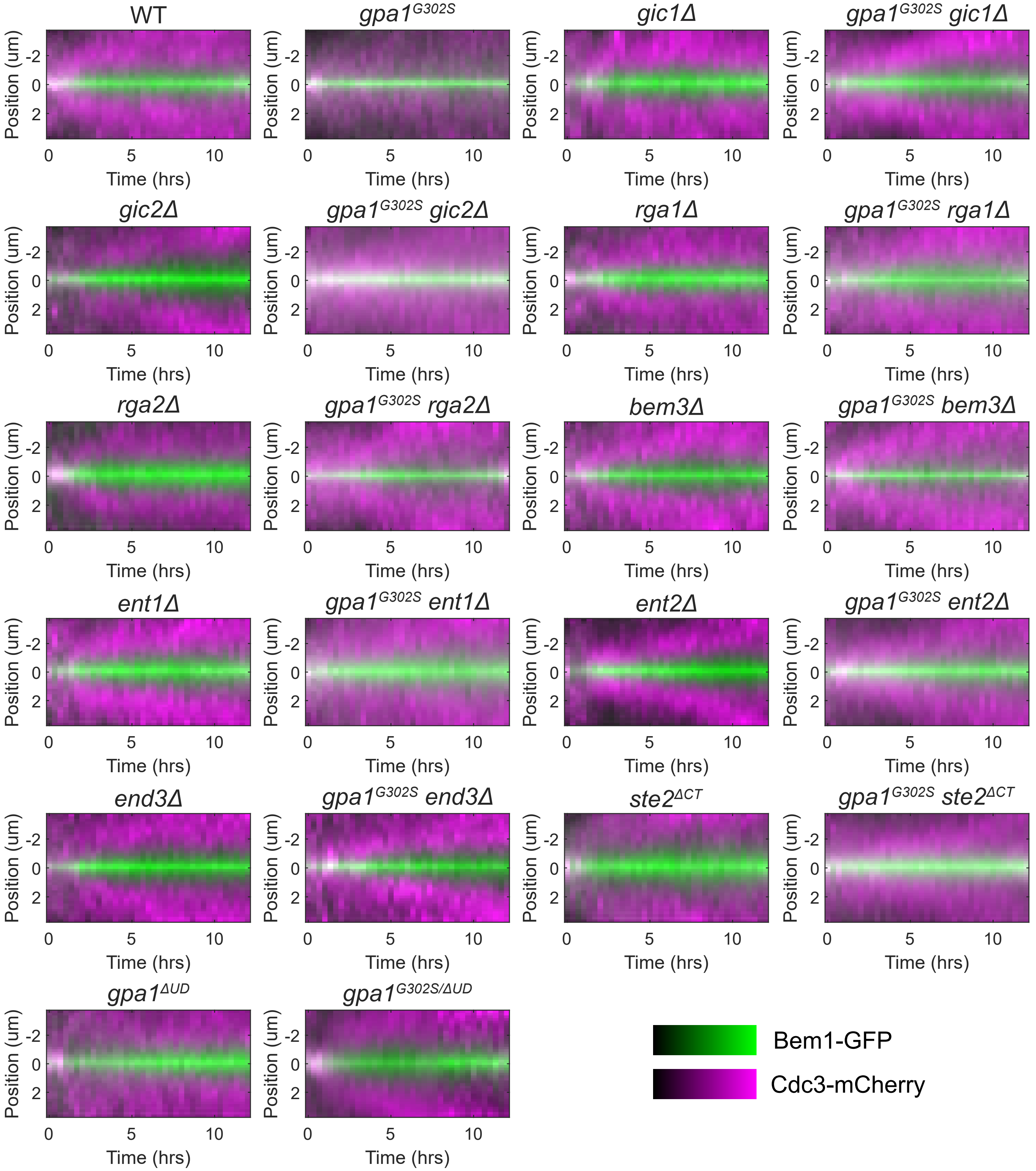


**Supplemental Figure S4. Merged Average Kymographs.** Merged images of the average kymographs of Cdc3-mCherry (magenta) and Bem1-GFP (green) from Supplemental Figures S2 and S3.

### Supplemental Figure S5

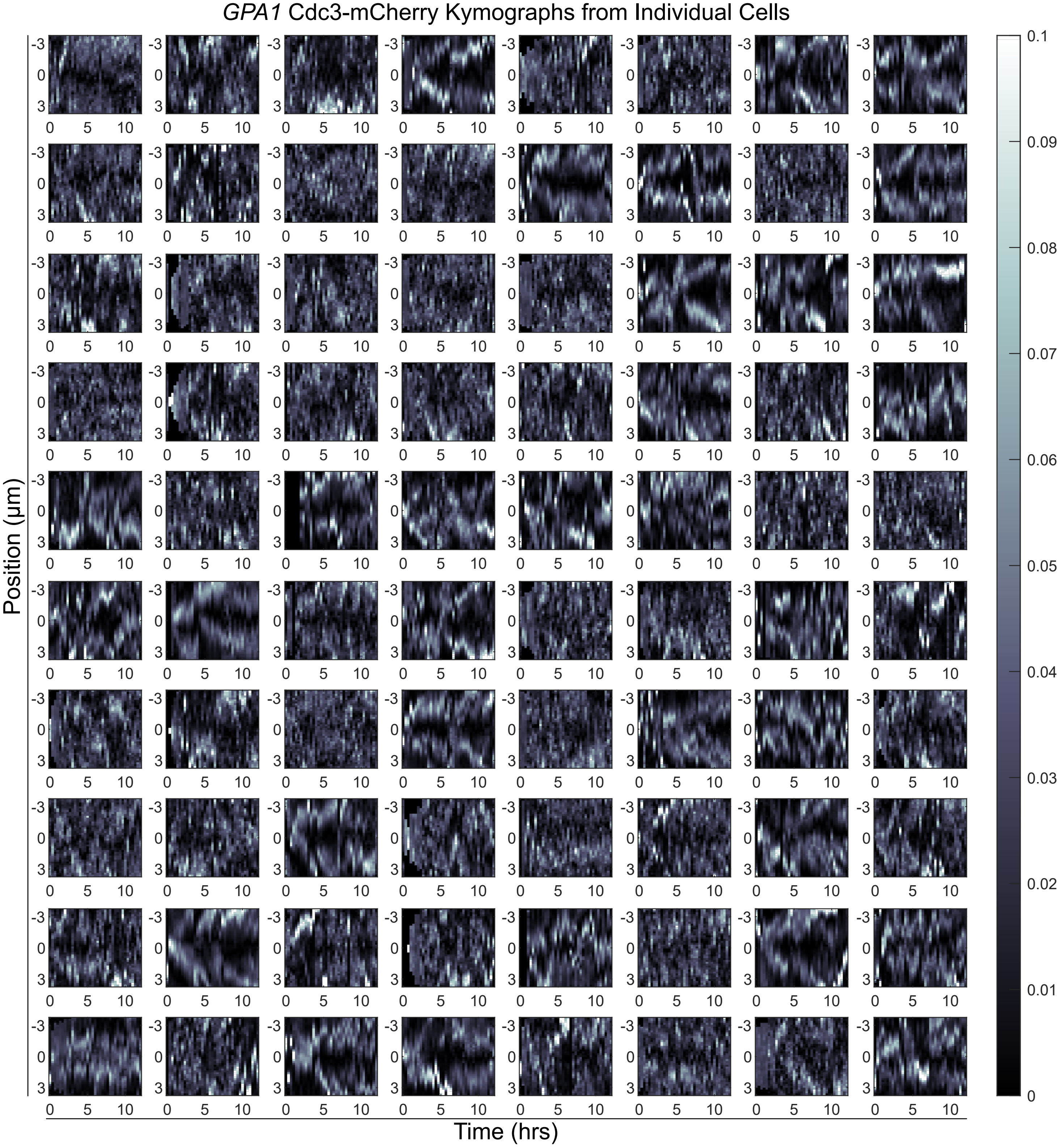


**Supplemental Figure S5. Individual Cdc3-mCherry Kymographs from WT cells.** Individual kymographs of Cdc3-mCherry from 80 individual WT GPA1 cells.

### Supplemental Figure S6

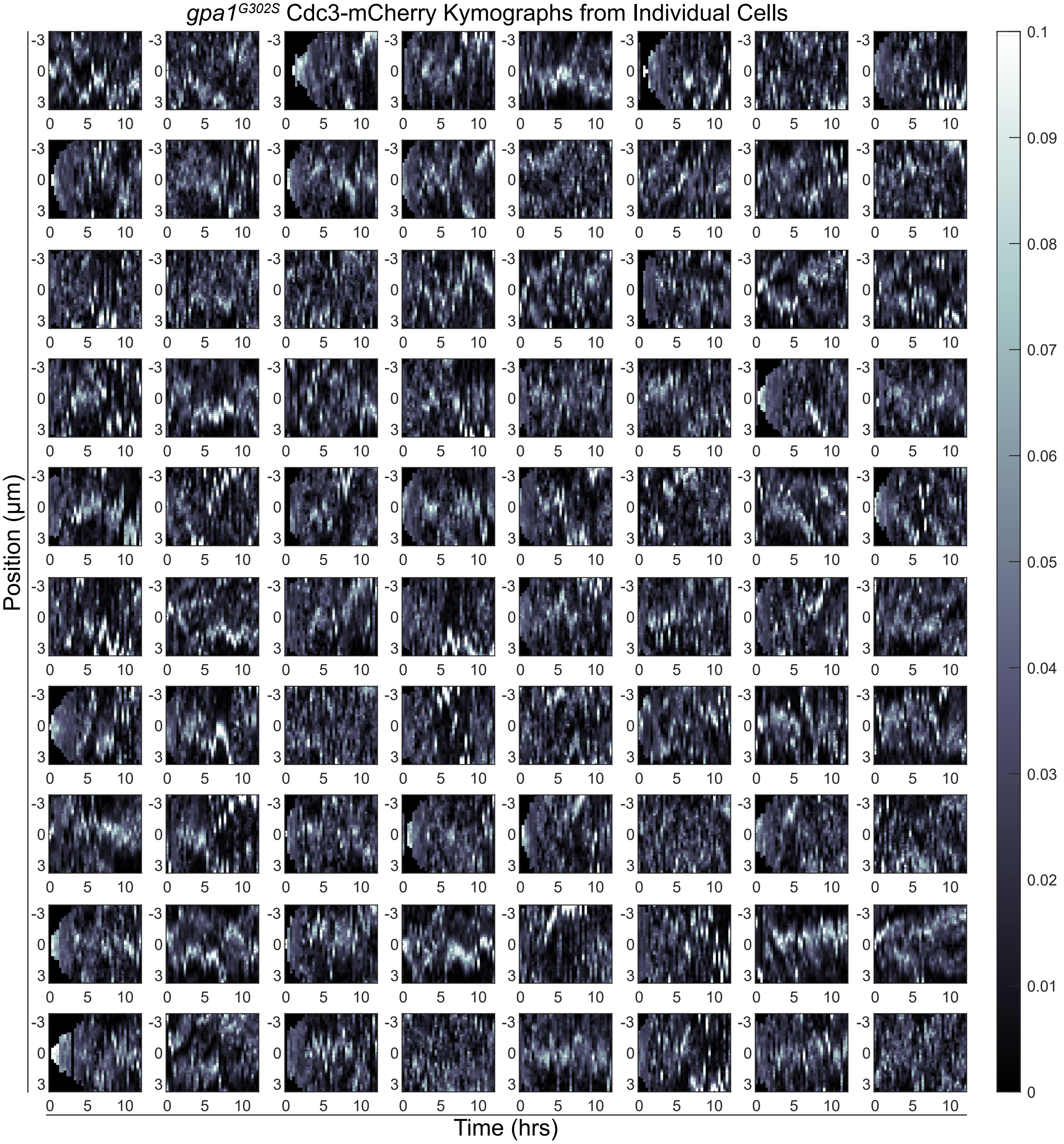


**Supplemental Figure S6. Individual Cdc3-mCherry Kymographs from gpa1$^{G302S}$ cells.** Individual kymographs of Cdc3-mCherry from 80 individual *gpa1$^{G302S}$* cells.

### Supplemental Figure S7

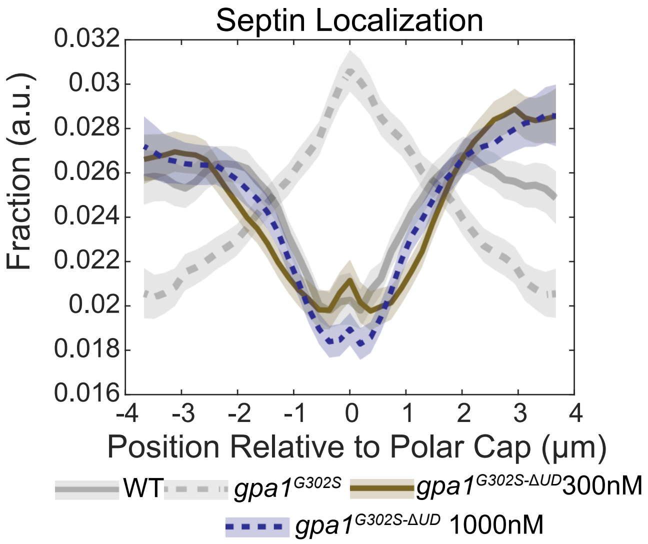


**Supplemental Figure S7. Altered sensitivity to pheromone in gpa1ΔUD cells does not impact quantitation of septin organization.** Quantitation of septin organization in cells expressing *gpa1*$^{G302S\text{-}\Delta UD}$ at 300 nM pheromone, conditions where they escape pheromone-induced arrest in G1, and at 1 μM pheromone, where cells stay arrested for the entire duration of the experiment.

### Supplemental Figure S8

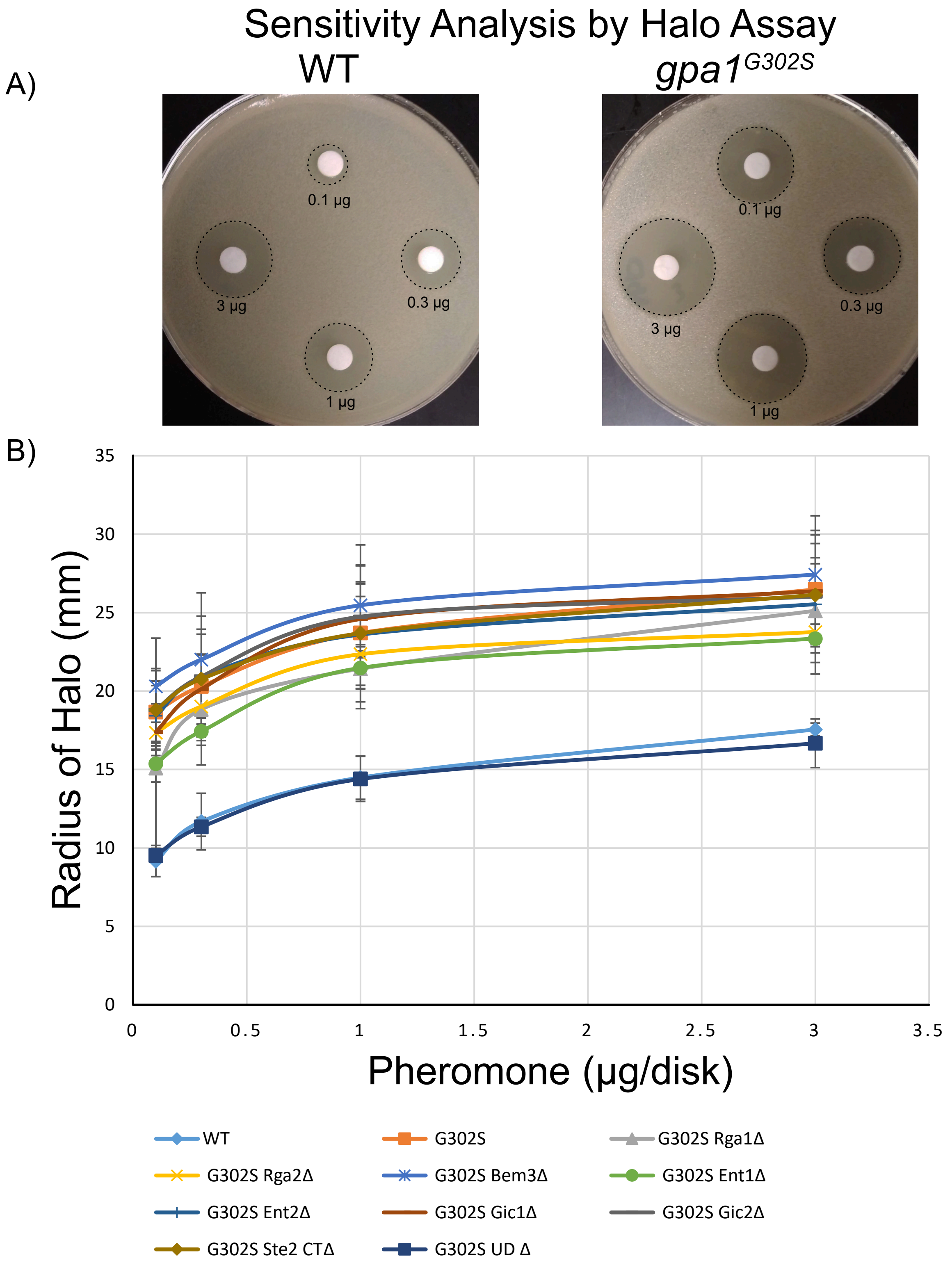


**Supplemental Figure S8. Sensitivity of *gpa1$^{G302S}$* background strains. A)** Halo assays were performed to assess the sensitivity of each strain to pheromone. Shown are examples of Halo Assays for WT and *gpa1$^{G302S}$*. Disks are labeled with the amount of pheromone present. B) Halo assays were quantified by measuring the halo radius (zone of no growth) around the indicated pheromone dose. Results were plotted as radius vs pheromone amount. Larger radii indicate higher sensitivity to pheromone. Each point is the average of 3 experiments. Error bars represent standard deviation.
